# Phylogeny, morphology, virulence, ecology, and host range of *Ordospora pajunii* (Ordosporidae), a microsporidian symbiont of *Daphnia* spp

**DOI:** 10.1101/2023.04.21.537887

**Authors:** Marcin K. Dziuba, Kristina M. McIntire, Kensuke Seto, Elizabeth S. Davenport, Mary A. Rogalski, Camden D. Gowler, Emma Baird, Megan Vaandrager, Cristian Huerta, Riley Jaye, Fiona E. Corcoran, Alicia Withrow, Steven Ahrendt, Asaf Salamov, Matt Nolan, Sravanthi Tejomurthula, Kerrie Barry, Igor V. Grigoriev, Timothy Y. James, Meghan A. Duffy

## Abstract

Impacts of microsporidia on host individuals are frequently subtle and can be context dependent. A key example of the latter comes from a recently discovered microsporidian symbiont of *Daphnia,* the net impact of which was found to shift from negative to positive based on environmental context. Given this, we hypothesized low baseline virulence of the microsporidian; here, we investigated the impact of infection on hosts in controlled conditions and the absence of other stressors. We also investigated its phylogenetic position, ecology and host range. The genetic data indicates that the symbiont is *Ordospora pajunii*, a newly described microsporidian parasite of *Daphnia*. We show that *O. pajunii* infection damages the gut, causing infected epithelial cells to lose microvilli and then rupture. The prevalence of this microsporidian could be high (up to 100% in the lab and 77% of adults in the field). Its overall virulence was low in most cases, but some genotypes suffered reduced survival and/or reproduction. Susceptibility and virulence were strongly host-genotype dependent. We found that North American *O. pajunii* were able to infect multiple *Daphnia* species, including the European species *D. longispina*, as well as *Ceriodaphnia spp*. Given the low, often undetectable virulence of this microsporidian, and potentially far reaching consequences of infections for the host when interacting with other pathogens or food, this *Daphnia* - *O. pajunii* symbiosis emerges as a valuable system for studying the mechanisms of context-dependent shifts between mutualism and parasitism, as well as for understanding how symbionts might alter host interactions with resources.

**Importance:** The net outcome of symbiosis depends on the costs and benefits to each partner. Those can be context dependent, driving the potential for an interaction to change between parasitism and mutualism. Understanding the baseline fitness impact in an interaction can help us understand those shifts; for an organism that is generally parasitic, it should be easier for it to become a mutualist if its baseline virulence is relatively low. Recently, a microsporidian was found to become beneficial to its *Daphnia* hosts in certain ecological contexts, but little was known about the symbiont (including its species identity). Here, we identify it as the microsporidium *Ordospora pajunii*. Despite the parasitic nature of microsporidia, we found *O. pajunii* to be, at most, mildly virulent; this helps explain why it can shift towards mutualism in certain ecological contexts and helps establish *O. pajunii* is a valuable model for investigating shifts along the mutualism-parasitism continuum.

## Introduction

Symbiosis can have profound impacts on the fitness of organisms. The outcomes of symbiosis span from being extremely positive and vital for both parties (e.g., plant pollination by insects, improved digestion in animals due to the gut microbiome) to being extremely detrimental to one of them (e.g., ebola virus is deadly to humans, *Batrachochytrium dendrobatidis* kills amphibians). Studies of microbial symbiosis have tended to focus on major groups of pathogens that cause diseases, particularly viruses, bacteria, and fungi. However, many groups of symbionts, such as microsporidia, are cryptic parasites that are unnoticed but can have important impacts on individual hosts or at the ecosystem level. Microsporidia are obligately intracellular and have highly reduced metabolism (1); they steal ATP and nucleotides from their host, inflicting negative consequences to host fitness (2). Therefore, they are (rightfully) considered parasitic (3). However, the impact of microsporidia can be context dependent, with them sometimes benefitting their hosts (4–6). This raises the question of how impactful microsporidia are to their hosts, and what determines when these obligate intracellular parasites could be beneficial?

Microsporidia can have a tremendous impact on host individuals as well as on populations, communities, and ecosystems. For example, the microsporidium *Paranosema locustae* has negative effects on the fitness of its locust hosts and also alters their behavior and disrupts swarm formation, which can positively affect crop survival (7). Microsporidia can alter entire communities even without harming the host; for example, *Microsporidia MB* inhibits the transmission of *Plasmodium*, the malaria causing agent, without negative consequences to the host mosquito (8), which, given the known effects of malaria on humans and animals, suggests this should have community-level effects. Moreover, some microsporidia, despite their parasitic nature, can provide more benefits than costs to a host; for example, the microsporidium *Nosema thompsoni* enables its native host ladybird to successfully invade new territories and outcompete local ladybird species by killing them (6).

The impacts of microsporidia are often mediated through virulence – that is, the damage that microsporidia cause to their hosts (e.g., (9, 10)). Microsporidian infections are frequently subclinical due to relatively low virulence, and the impact of microsporidia on their hosts can be context dependent (5, 11, 12). Whether the net outcome of a symbiotic interaction is positive (mutualism), neutral (commensalism) or negative (parasitism) is determined by the sum of the costs and benefits carried by the host (13). The costs of hosting a symbiont often stem from the necessity to share resources, but the magnitude of the cost is variable. Rapid host exploitation might increase the virulence of the symbiont, while more prudent exploitation could result in lower virulence (14). The sum of costs and benefits can be altered by abiotic and biotic factors (15), and while it is now clear that a symbiotic relationship can move along the mutualism-parasitism continuum, examples with well-characterized mechanisms underlying this shift are still scarce, particularly for typically parasitic organisms moving towards mutualism. To understand the mechanism underlying these shifts, we need to develop additional model systems where a comprehensive knowledge of symbiont biology and ecology allows for uncovering the costs and benefits of the interaction and how these change with ecological context.

A recent study found that infection of *Daphnia dentifera* (an ecologically important zooplankton host) with an unknown microsporidian can be beneficial for the host depending on the ecological context (4). The gut-infecting microsporidian was often found to have negative effects on *Daphnia*, especially when the host experienced food scarcity, though the relationship between food level and virulence varied between lakes (4). At the same time, infection with the microsporidian made the host’s gut less penetrable by spores of highly virulent fungus *Metschnikowia bicuspidata*. Intriguingly, *Daphnia* infected with the microsporidium had a reproductive advantage over the non-infected individuals during *M. bicuspidata* outbreaks. Overall, the net outcome of infection with the microsporidium spanned from parasitic through commensal to mutualistic, depending on the availability of food for the host and the prevalence of more virulent pathogens. Additionally, the effect size of infection with this microsporidian on the immune traits of *Daphnia* was found to be variable between lakes (16), which further indicates that the strength of the impact of the interaction on the host is context dependent. Thus, this *Daphnia*-microsporidium interaction has the potential to allow for better exploring the context dependency of symbiosis. However, while the host is a well-established model, with a wealth of information about its genetics, physiology and ecology (17, 18), little is known about the microsporidian. Indeed, in the prior studies showing context dependent outcomes (4, 16), it was referred to as “MicG” (because it is a microsporidian that attacks the gut), without a formal scientific name. Hence, the microsporidian needs further scrutiny under controlled laboratory settings to uncover its taxonomic identity and ecology and, most importantly, to quantify its virulence.

In this study we identified this *Daphnia*-infecting microsporidian previously called “MicG”, and then addressed the overarching question: what is the baseline cost of MicG infection? As the data on the ecology and especially virulence of this microsporidian is lacking, we investigated the consequences of infection on multiple clones of *Daphnia*, hypothesizing that the virulence of this pathogen would be relatively low to moderate, because that would make the shift from parasitism to mutualism that was observed in the field more likely. We also investigated the prevalence of this microsporidian in natural populations of *Daphnia*, hypothesizing that mild virulence and sometimes beneficial impacts of infection would support high levels of infection (19, 20). Together, this work not only establishes the identity of this intriguing symbiont, but also reveals more about its morphology, life history, and ecology.

## Materials and methods

### Overview

During an intensive study of host-symbiont interactions (4), we noticed a microsporidian infecting the guts of *Daphnia dentifera*. *Daphnia dentifera* Forbes is a dominant grazer in lake food webs in stratified lakes in the Midwestern United States (21) that hosts a variety of parasites (22, 23). The intensive study revealed that this microsporidian shifts along a mutualism-parasitism continuum (4) and spurred the work reported in this manuscript. The data on prevalence in four natural populations comes from the same field sampling as was reported in Rogalski et al. (4); however, that study did not analyze the temporal dynamics of infections.

### Sequencing and phylogenetic analysis

#### SSU rRNA sequencing

To investigate the phylogenetic affiliation of our microsporidian, we analyzed partial sequences of small subunit ribosomal RNA (SSU rRNA) of 9 individuals of *D. dentifera* infected with MicG collected in 2021 from Crooked Lake (“Crooked-W” in Table S1; Washtenaw County, Michigan, USA) and 9 individuals from Walsh Lake (Washtenaw County, Michigan, USA), against the sequences available in Genbank. DNA was extracted using DNeasy Blood and Tissue kits (Qiagen^lil^). Each infected *Daphnia* was picked from a microscope slide using jeweler’s forceps, placed in a 1.5 ml centrifuge tube with 100 µL and frozen until the DNA extraction was carried out. The defrosted animals were placed (with forceps) in 180 µL of ATL buffer, 20 µL of Proteinase K was added to each tube, and all tubes were incubated at 56°C for 24 hours. The next steps of the extraction were performed following the protocol described by the producer.

We amplified the SSU sequence of MicG using primers V1F (CACCAGGTTGATTCTGCCTGAC) and 530R (CCGCGGCKGCTGGCAC) (24) and KOD Xtreme™ Hot Start DNA Polymerase kit (Novagen^lil^). The PCR was performed using an initial step of 3 min at 95°C and then 35 cycles of 10 sec at 95°C followed by 30 sec at 60°C and 1 min at 72°C, and a final step of 5 min at 72°C. The PCR product was cleaned with ExoSAP-IT™ (ThermoFisher Scientific^lil^) kit using the manufacturer’s protocol, then Sanger sequenced by Eurofins Genomics^lil^ on an ABI 3730 XL Analyzer (Applied Biosystems) using forward and reverse primers on each product. The sequences obtained with forward and reverse primers were combined (after converting the reverse sequence into its reverse-complement version) and manually corrected to eliminate any sequencing errors. The MicG sequences were compared to sequences available in GenBank using blastn and the taxa indicated as the most similar were used to create a phylogenetic tree. The selected sequences were cropped to similar size and aligned using the ClustalW algorithm in MEGA X version 10.0.5 (25). A maximum likelihood phylogenetic tree was computed in MEGA X, using the Tamura 3-parameter model and 1000 bootstrap replicates.

#### ITS sequencing

Microsporidia are closely related to (or considered to be) fungi (26, 27), and the well-established molecular barcode for fungi is the Internal Transcribed Spacer (ITS) sequence (28). Therefore, we also analyzed ITS sequences of MicG infecting 7 *D. dentifera* from Crooked W Lake, 7 from Walsh Lake, and 1 *Daphnia mendotae* from Woodland Lake (Washtenaw County, Michigan, USA), all collected in 2021. The samples from Walsh and Crooked-W sequenced for ITS were a subset of the same samples as used for SSU (see above). We also analyzed ITS sequences of 8 *D. dentifera* from the ‘S’ genotype that had been infected in our lab culture of MicG isolated from Walsh Lake (isolate BDWalsh). The DNA extraction was performed using the same protocol as for SSU. The amplification was performed using primers ITSF-MicG (GGGGTGTGAGTCTTCTGTGG) and ITSR-MicG (GTTCAGCATCACACAACCCG) and the same PCR kit as above. The PCR parameters were an initial step of 5 min at 95°C and then 35 cycles of 30 sec at 95°C followed by 1 min at 53°C and 2 min at 72°C, and a final step of 5 min at 72°C. The product was purified with ExoSAP-IT™ and Sanger sequenced by Eurofins Genomics^lil^. The sequences were manually corrected and analyzed with the same software and methods as above, except for the alignment, which was performed with the Muscle algorithm and manually corrected.

#### Whole genome sequencing

We applied single cell genome sequencing techniques to sequence the MicG genome (29). A single *D. dentifera* was placed on a microscope slide in a droplet of MilliQ water and her gut was dissected, extruding the microsporidia. The spores were collected with a glass pasteur pipette and transferred into another droplet of MilliQ water, where large host tissue particles were removed. The spores were collected from the slide with a glass pipette and stored in a 1.5 mL centrifuge tube. Spores were isolated using a manually prepared drawn-out glass capillary pipette. A small amount (100–200 µL) of the spore suspension was transferred onto a petri dish and observed under an inverted microscope. We isolated ∼20 spores by pipette and put them into a small drop of UV-sterilized water. From them, 5–10 spores were isolated by avoiding putative contamination of host tissues and transferred into 200 µL PCR tubes with 1-2 µL of water. DNA extraction and whole genome amplification were performed using the Qiagen REPLI-g Single Cell kit (Qiagen, Maryland, USA). The prepared libraries were quantified using KAPA Biosystems’ next-generation sequencing library qPCR kit and run on a Roche LightCycler 480 real-time PCR instrument. Sequencing of the flowcell was performed on the Illumina NovaSeq sequencer using NovaSeq XP V1.5 reagent kits, S4 flowcell, following a 2x151 indexed run recipe. Ten million reads were filtered for JGI process contaminants with bbtools Seal, subsampled using <u>bbtools.reformat.sh</u> (http://sourceforge.net/projects/bbmap), and assembled using SPAdes version v3.15.3 (30), using parameters: --phred-offset 33 --cov-cutoff auto -t 16 -m 64 --careful -k 25,55,95. After the assembly, the scaffolds < 1000bp were removed. The genome was annotated using the JGI Annotation pipeline and shared via JGI MycoCosm (31).

Phylogenomic analysis was performed using a dataset of 40 Microsporidia taxa (Table S2). Two species of Metchnikovellidae (*Amphyamblys* sp. and *Metchnikovella incurvata*) were selected as the outgroup. The “genome mode” of BUSCO version 5 (32) was run for each microsporidian genome using the microsporidia_odb10 database. A concatenated alignment was prepared using the version 4 of “BUSCO_phylogenomics” pipeline (https://github.com/jamiemcg/BUSCO_phylogenomics (33)). Single copy BUSCO genes present in more than 80% of 40 microsporidians (“-psc 80”) were extracted from the outputs of BUSCO. Each gene sequence was aligned with muscle v3 (34) and trimmed with trimAl v1.2 (35). The final concatenated alignment comprised 102,302 amino acids from 332 genes. A maximum likelihood (ML) analysis was performed using IQ-TREE v2 (36) using the site-heterogeneous model (LG + C60 + F + G + PMSF (37)). The tree inferred using the LG + F + R9 model selected by ModelFinder (38) was used as a guide tree for PMSF model analysis. A standard nonparametric bootstrap analysis (100 replicates) was done under the LG + C60 + F + G + PMSF model.

Based on phylogenetic similarity to *Ordospora pajunii* FI-F-10, we performed comparative analyses to determine the extent of genetic differences between the genomes of *O. pajunii* and MicG. The genomes were first compared to calculate average nucleotide identity using OAT 0.93.1 with the OrthoANI algorithm (39). Alignment of FI-F-10 and MicG was conducted using Parsnp v1.2 (40) to identify single nucleotide polymorphisms (SNPs). We used SnpEff 5.0e (41) to calculate the putative functional effects of the SNPs using the annotation of FI-F-10 as a reference. To facilitate comparison to FI-F-10 we first annotated the genome using PRODIGAL v2.6.3 (42), and the two annotations were compared using OrthoFinder v. 2.5.2 (43) to identify genes that were missing or duplicated in one or the other genome assemblies. A final genome annotation was performed using the JGI pipeline (31), and the genome is available at https://mycocosm.jgi.doe.gov/MicG_I_3.

### Microscopy

We used a combination of light and transmission electron microscopy to explore the morphology of MicG (isolate BDWalsh), as well as the symptoms associated with infection.

#### Light microscopy

We used an Olympus DP73 camera attached to an Olympus SZX16 dissecting microscope to take photomicrographs of *D. dentifera* infected with MicG at 4 - 11.5x magnification. In addition, MicG spores were extracted by rupturing the gut of four infected *Daphnia* on a microscope slide; the length and width of 25 spores from each *Daphnia* were measured using the same camera attached to an Olympus BX53 microscope (1000x magnification) and using Olympus cellSens Imaging Software.

#### Transmission electron microscopy

To characterize the ultrastructure of MicG, as well as the impact on infected host cells and gut morphology, we collected twenty MicG-infected *D. dentifera* individuals from lab cultures (see Supplemental Materials), fixed them in 3% glutaraldehyde, and stored at 4°C for over two weeks. After primary fixation, samples were washed with 0.1M phosphate buffer and postfixed with 1% osmium tetroxide in 0.1M phosphate buffer, dehydrated in a gradient series of acetone and infiltrated and embedded in Spurr’s resin. 70 nm thin sections were obtained with a Power Tome Ultramicrotome (RMC, Boeckeler Instruments, Tucson, AZ) and post stained with uranyl acetate and lead citrate. Images were taken with a JEOL 1400Flash Transmission Electron Microscope (Japan Electron Optics Laboratory, Japan) at an accelerating voltage of 100kV.

### Life table measures of parasite virulence

To quantify parasite virulence (with respect to reproduction and lifespan), we carried out life table experiments with 7 genotypes of *Daphnia dentifera* (S and A43, originally isolated from lakes in Barry County, Michigan, and M37, ML32, BD08, DW29, and DW22, originally isolated from lakes in Sullivan and Greene Counties, Indiana; the ‘S’ clone is also referred to as “Standard” or “Std” in some other publications). Each of those genotypes is a clonal line that has been kept for years in the laboratory. We first reared individuals from these genotypes for 3 generations in standard conditions to standardize maternal effects. We used two different exposure times in our experiment. Our preliminary work on this parasite had suggested that younger hosts might be more susceptible, and there is substantial evidence that age can influence infection outcomes (44, 45).

On experiment day 1, 0-24h old neonates were collected and placed individually into 50mL beakers, with 30mL filtered lake water and 1mL standard food stock (to a final concentration of ∼33,000 cells/mL of *Ankistrodesmus falcatus*). Animals were assigned to one of three pathogen treatments: MicG 48h, receiving one dose of MicG at 48h old and a placebo at 120h old; MicG 120h, receiving a placebo at 48h old and one dose of MicG at 120h old; and Control, receiving a placebo at 48h and 120h. Two exposure windows were chosen as previous observations suggested MicG may more effectively infect hosts very early in the host lifespan. There were 10 replicate individuals per clone * exposure treatment combination (210 individuals total). Animals were housed in environmental chambers at 20°C with a 16h:8h (light:dark) cycle for the duration of the experiment.

On day 2, all animals were transferred to new beakers with 10mL filtered lake water and given either a dose of MicG isolate BDWalsh or a placebo. For MicG doses, a slurry was created by crushing *Daphnia dentifera* (S clone, in which we cultivate this symbiont) with high MicG infectious burden into 10mL MilliQ water; this slurry contained a ratio of 1 high burden donor: 1 experimental animal, with each experimental animal exposed to a portion of the slurry equivalent to 1 high burden donor (e.g., if 10 animals were being exposed, the slurry would be created using 10 highly infected donors, and each experimental animal would receive 1/10th of the resulting slurry). The placebo was created and dosed in the same manner, using uninfected animals. After 48 hours, animals were moved to new, parasite-free beakers with 30mL filtered lake water. On day 5, the exposure treatment was repeated as above; this time, animals in the MicG 120h treatment received the spore slurry and the other two treatments received the placebo. During exposure, animals were given 0.5mL stock food per day, all other experimental days animals received 1mL stock food daily (1,000,000 *A. falcatus* cells/mL in the stock food solution).

We checked animals daily for mortality and offspring production. We recorded the date and number of offspring produced (offspring were removed from the beaker) as well as the date of death. Animals were checked for infection twice per week under the dissecting microscope (4- 10x magnification).

We calculated the proportion of exposed hosts that became infected for each combination of genotype and infection timing. We then tested for effects of the parasite and timing of parasite exposure on host life history traits. We used an aligned rank ANOVA to analyze total fecundity, a generalized linear model (GLM) to analyze body size at first reproduction, and a proportional hazard model by genotype to analyze time to death. These analyses were restricted to the four genotypes that displayed substantial infection (BD08, DW29, DW22 and S). The analyses included all individuals in a genotype*treatment combination; results were not qualitatively different if only infected individuals were included for the MicG 48h and MicG 120h treatments.

### Prevalence in natural populations

We sampled four lake populations in Southeastern Michigan in 2016: Gosling Lake in Livingston County and North, Pickerel, and Sullivan Lakes in Washtenaw County (Table S1). These lakes were sampled every three days from late July through mid-November. This sampling window was chosen because outbreaks of other *D. dentifera* parasites occur primarily in late summer and autumn (46, 47). Each lake was sampled by taking whole water column tows with a Wisconsin net from three locations in the deep basin of the lake. These three tows were pooled, and then a subsample was analyzed to determine the prevalence of individuals that were infected; this was done by examining each individual in the subsample under a dissecting microscope at 40-50x using dark field microscopy, looking for characteristic signs of infection along the gut (described more below). We aimed to have at least 200 *D. dentifera* individuals in the subsample that was analyzed for infections; if there were fewer than 200 individuals in the whole sample, we analyzed the entire sample. Over the entire study period, the median number of *D. dentifera* that we analyzed for infection was 233.5 per lake-day combination (mean: 236, minimum: 71, maximum: 449). Because adults are more likely to have visible signs of infection, we focus on the proportion of the adult population that was infected in our analysis. We plotted the raw data and fitted a loess curve with a span of 0.4 for each lake.

### Host range

We used a dataset from 15 inland lakes in Southeastern Michigan (Table S1) collected in 2021 to assess which *Daphnia* species are infected by MicG in natural populations. The 15 lakes are the same as those in Gowler et al. (48). All of these lakes are stratified and include *D. dentifera* as a dominant member of the zooplankton. We also analyzed one sample from Lake Erie.

To check whether MicG can infect European *D. longispina* (the host from which *O. pajunii* was described (49)), we exposed five clones of *D. longispina* collected from three German lakes, along with the *D. dentifera* clone S (used as a control for infection success) to MicG isolate BDWalsh spores. We synchronized the mothers of experimental animals, and reared them until they gave birth to their first clutch. The first clutch babies were harvested within 24h of birth and placed singly in a 50mL beaker filled with 20mL of filtered lake water and fed 0.5mL of a standard food stock (1,000,000 *A. falcatus* cells/mL in the stock food solution) daily until the end of exposure. We exposed 10 individuals from each of the 6 clones. After 48h, all animals were dosed with a MicG spore slurry; each animal was exposed to a spore suspension equivalent to one donor individual, similar to the approach used in the life table experiment above. Exposure lasted 48h, after which we transferred *Daphnia* to new beakers filled with 40mL of filtered lake water, fed them 1 mL of stock food daily and checked for mortality until the end of the experiment. We initially checked *Daphnia* for infection under the dissecting microscope 7 days after exposure, then checked them twice a week thereafter. The experiment ended on day 30. The S clone individuals that we used as a positive control had 100% MicG infection prevalence in this trial.

## Results

### Sequencing and phylogenetic analysis

The SSU and ITS sequences of MicG suggest a very close affinity with *Ordospora pajunii* (Figure 1), a microsporidian species recently described from *Daphnia longispina* in Finland (49). Moreover, there was no variation within the MicG isolates at SSU or ITS (Figure 1A, B); this supports that our diagnostic criteria (based on visible symptoms) identify a single species. It is also notable that the isolates from our 2021 collections had identical SSU sequences to the sample collected in 2016 as part of the study exploring shifts along a mutualism-parasitism gradient (4) (accession number: MH635259.1; Figure 1A). There was very little variation between MicG and the published sequences of *O. pajunii* (Figure 1A, B). The average nucleotide identity between MicG and *O. pajunii* calculated for SSU sequence was 100%. There was a clear separation from the *Ordospora colligata* clade, however, at both loci.

**Figure 1.**
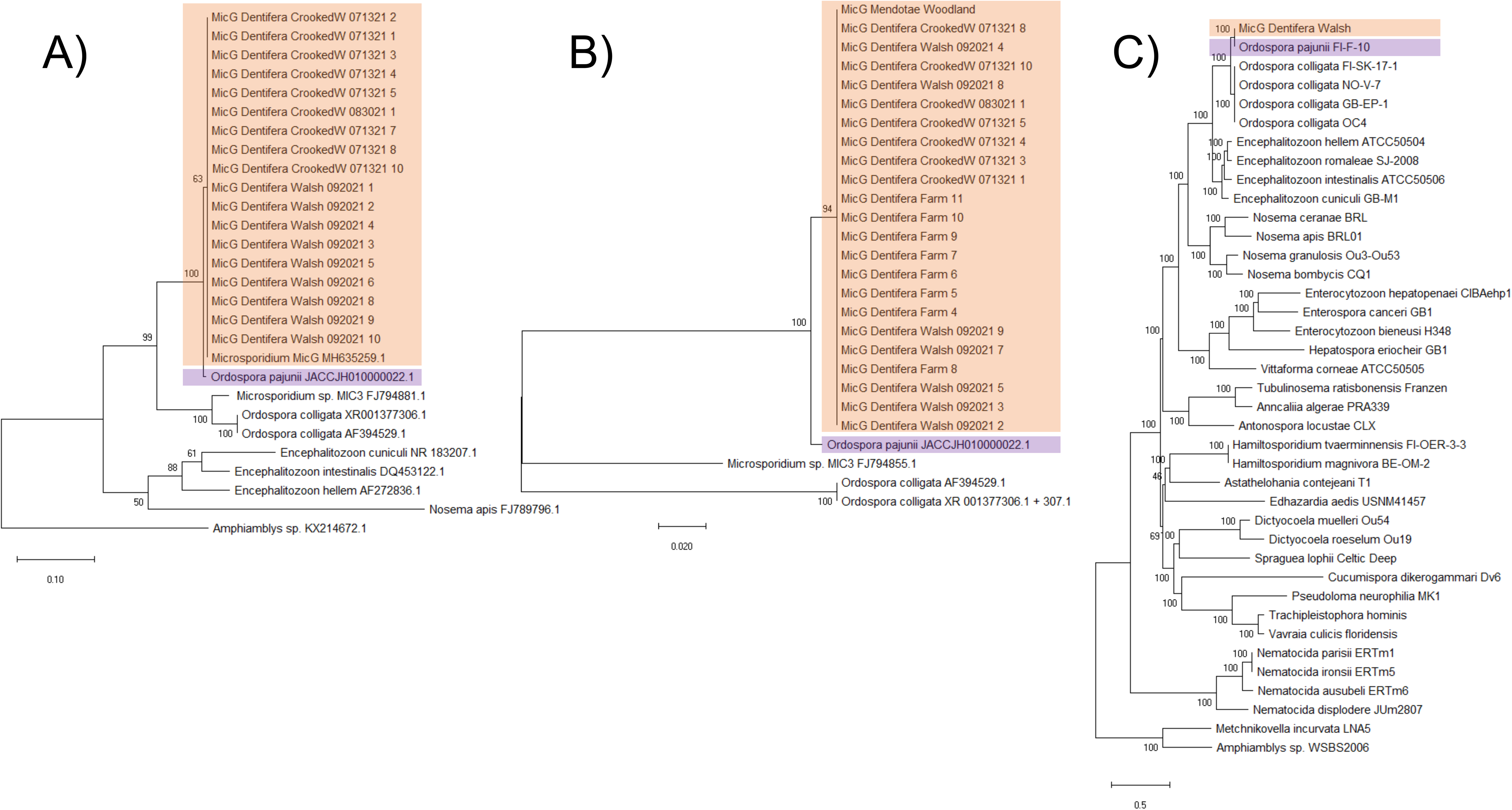
Phylogenetic analyses suggest a very close affinity of MicG and the recently described *O. pajunii*. A) A maximum likelihood phylogenetic tree of partial SSU ribosomal RNA sequences reveals no variation within MicG, and high similarity to *O. pajunii*. The sequence labeled “Microsporidium sp. MicG MH635259.1’’ was previously sequenced as part of the Rogalski et al. (4) study. B) ITS sequences of MicG were also identical and very similar to that of *O. pajunii*. In A&B, the branch support is based on 1000x bootstrap replicates, and the scale corresponds to the branch length and represents the number of substitutions per site. C) A maximum likelihood phylogenetic tree of 332 genes from the microsporidia_odb10 database also supports very high similarity between MicG and *O. pajunii*; branch support is based on 100x bootstrap replicates. MicG isolated from lakes in Michigan, USA and *O. pajunii* from Finland (49) are indicated with orange and purple rectangles, respectively.

We assembled the MicG genome into 42 contigs of total length 2.24 Mb with a contig n50 of 133 kb. A phylogenomic analysis using the amino acid sequences of 332 genes corroborated the high genetic similarity of MicG to *O. pajunii* (Figure 1C). The average nucleotide identity between MicG and *O. pajunii* was 99.84%, which is well within the 95% limit recently suggested for species (49). The MicG genome annotation revealed 1946 protein coding genes. Alignment of the MicG and *O. pajunii* genomes uncovered 2,666 SNPs and 1,004 bp associated with indels differing between the strains. 12.8% of the SNPs were intergenic, and, of the SNPs in exons, 55.3% were missense, and 8 total nonsense SNPs were observed. Comparison of the two largest contigs of MicG to *O. pajunii* revealed colinearity of chromosomes (Figure 2), with most of the variation between the two genomes appearing at the ends of contigs. High variability of the genome in the subtelomeric regions with associated repeat regions and a ribosomal RNA gene array is also observed for other microsporidia (50). Ribosomal RNA, however, was not associated with the ends of contigs in *O. pajunii* or MicG and mostly absent from the genome assemblies save one small contig in each genome, suggesting our assemblies are incomplete towards the telomeres. Beyond the subtelomeric regions, we inspected genes with the highest number of amino acid changing SNPs between the strains. The highest number of missense mutations (Figure S1) were in: an alpha/beta hydrolase (20 SNPs; *O. pajuni* protein accession number: KAH9410807.1), a hypothetical protein (11 SNPs; KAH9411254.1), a DEAD/DEAH box helicase (7 SNPs; KAH9411242.1), and an aminopeptidase (7 SNPs; KAH9410533.1). There was no evidence for genes present in one strain but not the other when only considering the large contigs (>10 kb).

**Figure 2.**
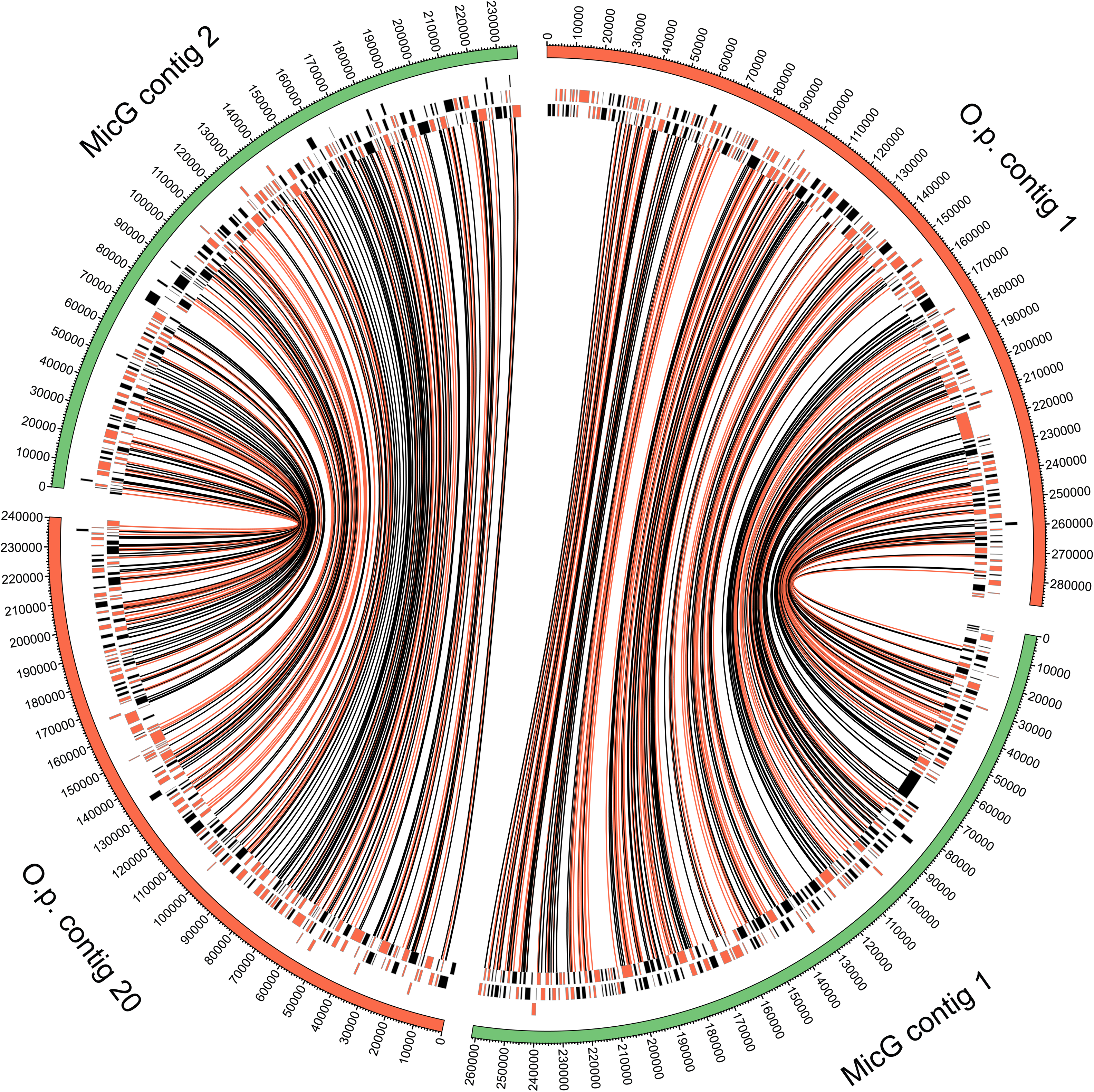
Comparison of two largest contigs of MicG with homologous contigs of *O. pajunii* shows conserved gene order. The outer band shows the lengths of chromosomes, and the inner rings are the locations of the genes, arbitrarily split into three lines to prevent overlap for clarity (black markers are on + strand and orange markers on - strand). Links connect SNPs identified following alignment and are indicated as amino acid changing (orange) or silent (black). Plot generated using Circos (51).

The high similarity across the entire genomes of MicG and *O. pajunii*, as well as complete concordance of sequences at the SSU region and minimal difference at the ITS locus between MicG and *O. pajunii* from Europe, lead us to conclude that they are the same species. Nonetheless, we found some differences between North American and European *O. pajunii*. We identified genetic divergence in some of the coding regions – most notably a 30 bp indel in the *chitin synthase* gene. The sequence of the MicG *chitin synthase* gene in this region is almost identical to that of *O. colligata* (Figure S2), suggesting a deletion in European *O. pajunii*. We also identified multiple missense and nonsense mutations in other genes when comparing the genome of MicG with *O. pajunii*.

### Microscopy

We observed the first symptoms of infection about 7-10 days after exposure. Infection is visible as white globular structures embedded within the host’s gut epithelium (Figure 3A, B). At first, only several clusters of the parasite spores are visible. Spores initially occupy the central part of the host’s cell (Figure 3G), but as they grow and divide, the vacuole containing them starts expanding and growing towards the inside of the host’s gut lumen (Figure 3H). At the late stage of cluster development, the cluster is visible as a bubble-shaped sac (Figure 3C, D) waved by the peristalsis of the host’s gut. The vacuole eventually bursts, releasing the spores into the gut lumen (Figure 3C, I). The spores are then shed into the environment, although it is possible (perhaps even likely) that some reinfect the same host prior to release into the environment. Interestingly, electron microscopy revealed that the microvilli are absent from the epithelial cells in these later stages of cluster development, when the cluster has moved towards (or into) the gut lumen (Figure 3F, H, I).

**Figure 3.**
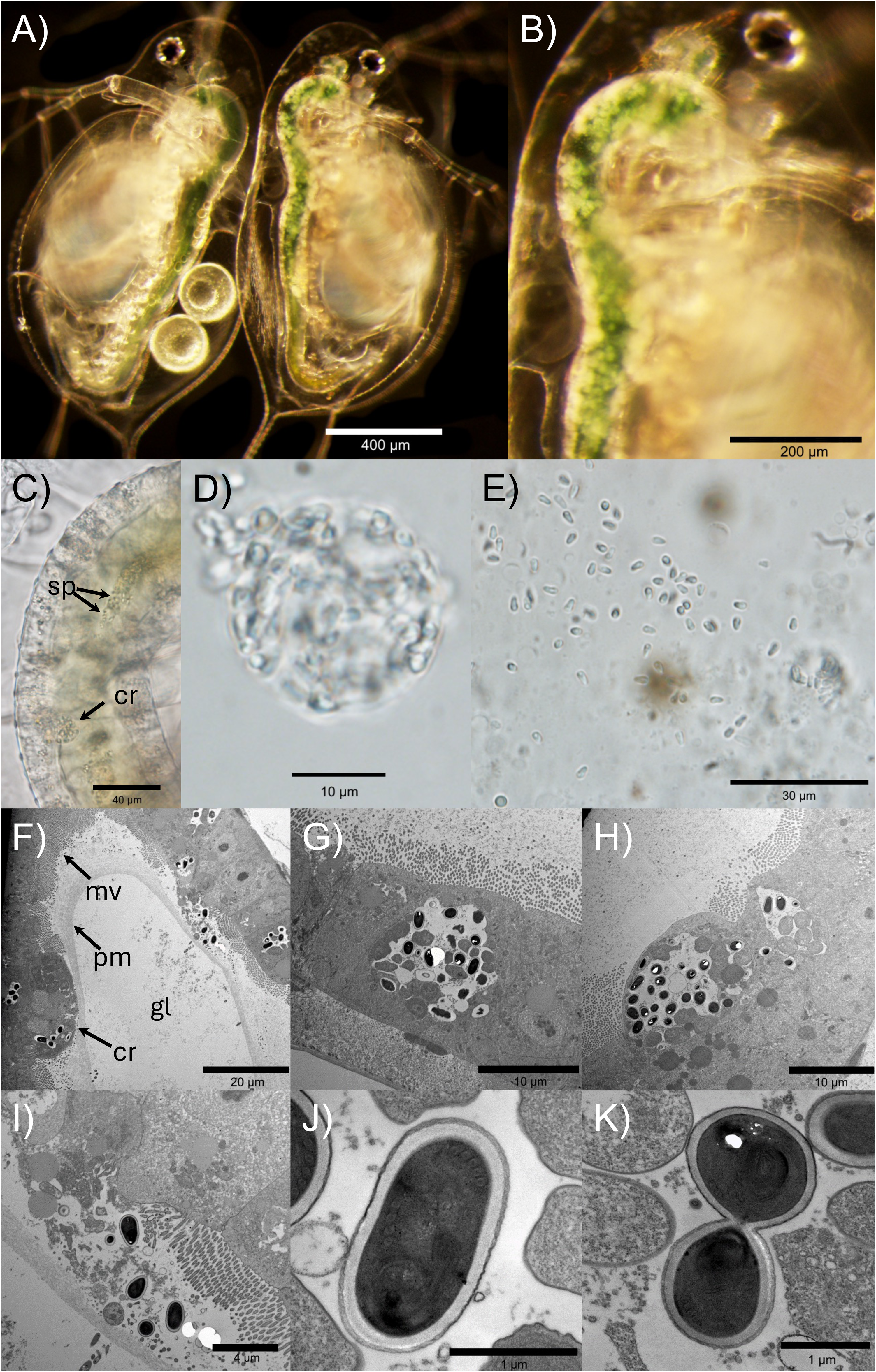
A) *D. dentifera* infected with *O. pajunii* (on the right) vs. uninfected *Daphnia* (on the left); note that white opaque spore clusters fill the gut of the right *Daphnia*, but not the left one. The infection is most obvious in the upper half of the gut. The globular but more transparent and larger structures on the dorsal middle part of the left (uninfected) *Daphnia* gut are not the parasite spores; instead, these are fat cells and are outside of the gut lumen, B) the same infected *Daphnia* as in ‘A’, at higher magnification, C) upper part of midgut of an infected *Daphnia*, with a mature spore cluster (cr) and visible spores coming from a burst cluster (sp), D) a cluster of *O. pajunii* spores extracted from the gut, E) free floating spores extruded from the gut; after extracting a host’s gut, the clusters rapidly burst and release the spores, F) cross section of an infected host’s gut with visible microvilli (mv), peritrophic membrane (pm), a maturing spore cluster (cr) and gut lumen (gl) indicated, G) a spore cluster in a central part of the epithelial cell, H) a more advanced spore cluster growing towards the inside of the gut lumen, I) spores released from the host’s cell after a cluster burst, J) a single spore of *O. pajunii*, and K) a late stage of cell division. Pictures A-E were taken using light microscopy, while pictures F-K are taken with transmission electron microscopy (TEM); the latter involves staining which makes the spores dark. The photos show MicG isolate ‘BDWalsh’ infecting the ‘S’ genotype, with the animals collected from MicG ‘farms’ in the lab.

Initial infections most frequently occur in the upper part of the midgut and diverticula. Over time, the number of clusters and general coverage of the gut by the parasite increases, resulting in tens to hundreds of clusters spread across the gut, frequently merging into an indistinguishable mass (Figure 3A, B). The spores are about 3.03 µm long (range: 2.57-3.71, n=100) and 1.66 µm wide (range: 1.45-1.96, n=100), initially oval (Figure 3J, K) and pyriform when mature (Figure 3D, E). The size of MicG spores overlaps with *O. colligata* and the *O. pajunii* isolate described by de Albuquerque et al. (49); however, unlike those, MicG does not form chains when extruded from the host (Figure 3E).

### Life table measures of parasite virulence

Infection prevalence varied across genotypes from 0-100% infected individuals (Figure 4A). Early exposure, as opposed to late exposure, to parasite spores resulted in similar or greater infection success among the susceptible clones (Figure 4A). Additionally, we observed consistent variation in the morphology of infection across genotypes (Figure 4B). Infection in the S clone manifests as densely filled globules with defined edges; in BD08, infection frequently manifests with a grainy appearance where the clusters are much smaller and more dispersed; finally, in DW22, infection manifests as an amorphous cloudy patch in the gut.

**Figure 4.**
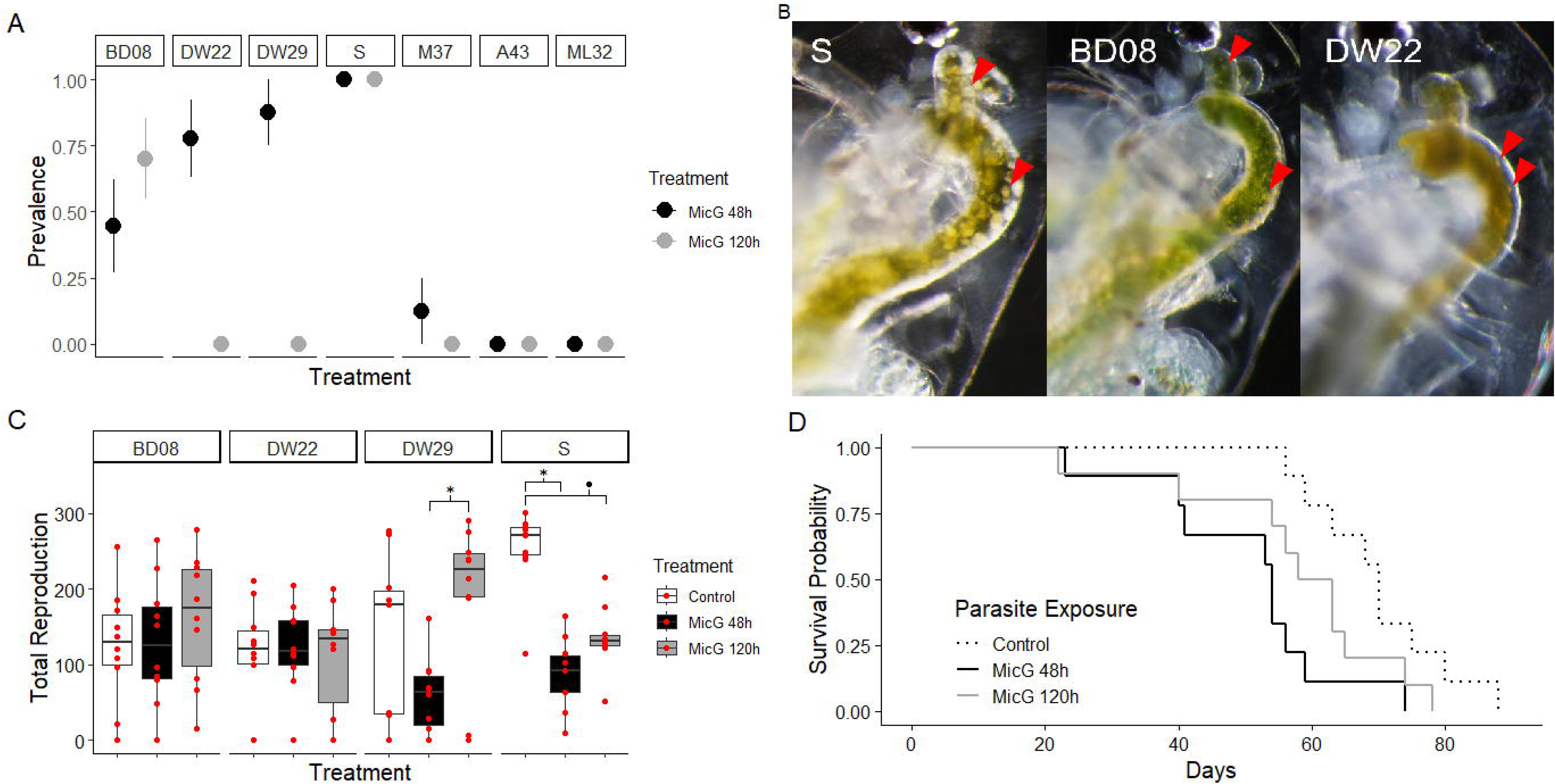
Genotypes differed in their likelihood of becoming infected, in the morphology of infection in the gut, and in the impact of infection. A) Clones varied in the proportion of animals displaying visible infection (clone name indicated on top). The point and error bar represents the binomial mean and standard error. B) Morphology of infection with MicG in three of the susceptible clones: S, BD08, DW22. The red arrows indicate examples of MicG clusters. C) The parasite impacts lifetime total reproduction (that is, the total number of offspring produced across the experiment) in genotypes differently. The horizontal line represents a median, the boxes show first and third quartile and the whiskers represent the 1.5*IQR from the top and bottom. Raw data are overlaid in red. *p<0.05, ·p<0.1 D) Time to event analysis for mortality indicated that only the lifespan of the S genotype was impacted by MicG (for other clones see Fig. S4). S genotype lifespan was significantly shortened with early exposure (MicG 48h, solid black line) compared to no parasite exposure (Control, dashed line; hazard ratio=4.577, 95%CI=1.616-12.964, P=0.0042).

Total lifetime fecundity varied depending on parasite treatment and clone (interaction: F_6,108_=5.945, P<0.0001). More specifically, the exposed S genotype produced fewer offspring than the controls if they had been exposed early in life (MicG 48h treatment vs. control: T_108_=4.908, P=0.0002) and marginally fewer if they were exposed later (MicG 120h treatment vs. control: T_108_=3.297, P=0.076, Figure 4C). In addition, the DW29 genotype differed between parasite treatments, with early exposure animals producing fewer offspring compared to later exposure MicG 48h vs. MicG 120h: T_108_=4.082, p=0.006, Figure 4C). Host body size did not differ depending on parasite treatment (F_2,69_= 0.1482, P=0.8626, Figure S3). Host time to death differed due to parasite treatment, but only for the S genotype; the lifespan of this genotype was significantly shorter in the early parasite exposure compared to the no parasite treatment (hazard ratio=4.577, 95%CI=1.616-12.964, P=0.0042, Figure 4D, Figure S4).

### Parasite prevalence in natural populations

Two of the four lakes, North and Pickerel, had sustained outbreaks of MicG (Figure 5). North Lake already had ∼20% of *D. dentifera* adults infected at the beginning of the sampling period (in late July and early August). In North Lake, infection prevalence reached its maximum in mid-September (48% of the total *D. dentifera* population, 77% of adult females); in Pickerel, infection prevalence reached its maximum in the first half of October (57% of the total *D. dentifera* population, 70.5% of adult females). Infections were much lower in the other two lakes, except for a spike in infections in Sullivan near the end of the sampling period (in November); the population size remained high through this time, so this increase is not an artifact of low sample sizes (Figure S5).

**Figure 5.**
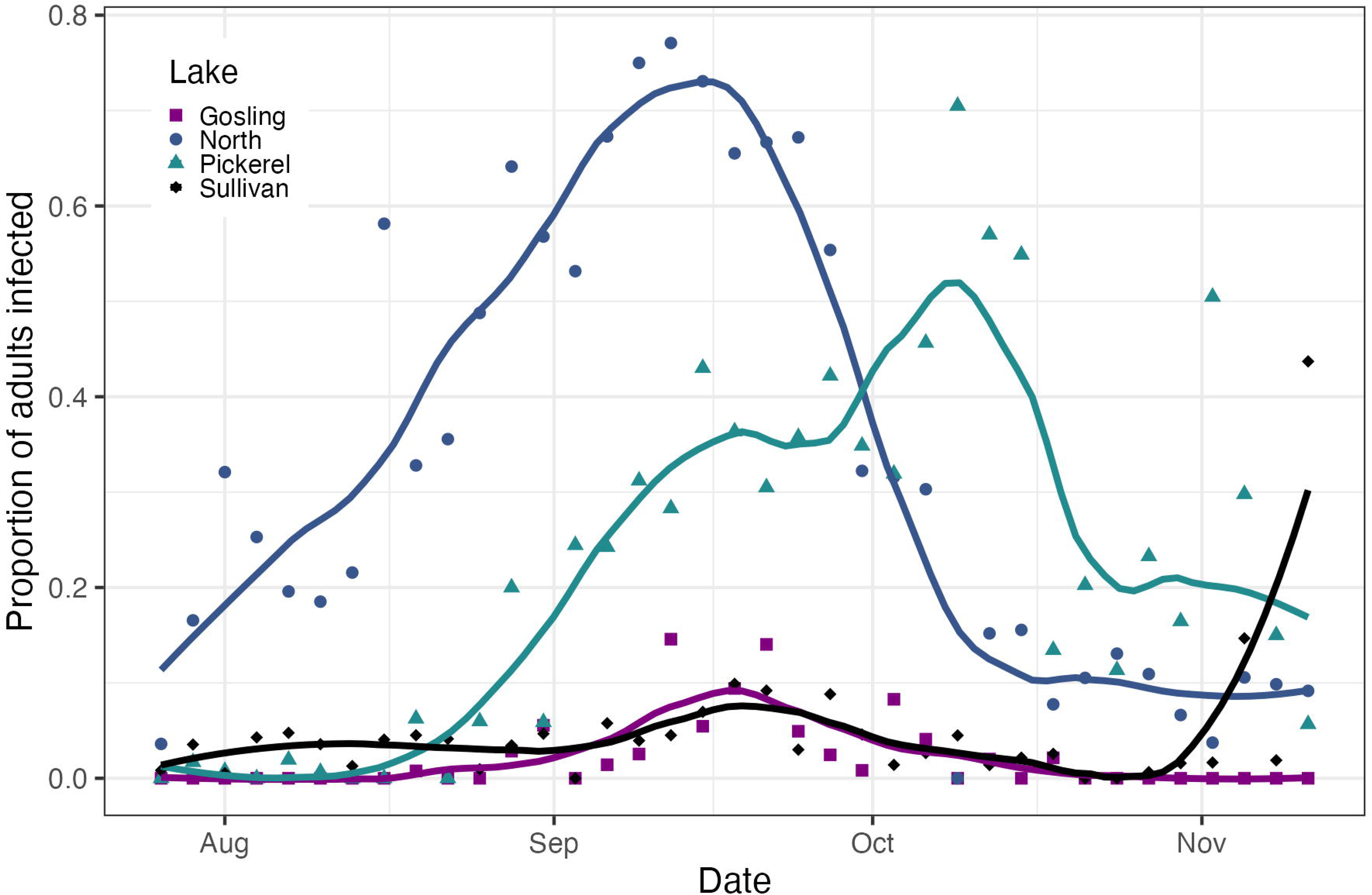
The proportion of *D. dentifera* adults that were infected with MicG varied over time and across lakes. Data come from four lakes (Gosling, North, Pickerel, and Sullivan) in Southeast Michigan, USA, that were sampled every three days in 2016. Raw data and fitted loess curves are presented.

### Host range

We observed MicG infections in *Ceriodaphnia dubia*, *Daphnia ambigua*, *D. dentifera*, *D. dubia*, *D. mendotae*, *D*. *parvula*, *D. pulicaria* and *D. retrocurva* – that is, all of the *Daphnia* and *Ceriodaphnia* species in our study lakes. In addition, in our laboratory infection assay, MicG infected two out of the five *D. longispina* clones. Similar to *D. dentifera*, exposed *D. longispina* clones varied in their susceptibility to MicG (spanning 0% to 90%, Table S3).

## Discussion

Prior ecological studies on this microsporidian have shown a clear impact on its host and on other symbionts competing for the same host (4, 16), as well as shifts along a mutualism-parasitism continuum; this all suggests promise as an interesting study system, especially since the host is already a model organism (17, 18, 52). In the present study, we characterized the phylogeny, morphology, virulence, ecology, and host range of this symbiont, with a goal of understanding the impact of this pathogen on the host and facilitating future work on this system. Our phylogenetic analyses indicate very high similarity between MicG and *Ordospora pajunii* (Figure 1 and Figure 2), a microsporidian recently described from *Daphnia longispina* in Europe (49). The spore morphology of both microsporidians is alike and both infections have similar presentation in the host’s gut, overgrowing the epithelium and being successively shed and excreted with feces. We found that MicG infections result in loss of microvilli of the infected cells, which likely affects the host’s interaction with resources. MicG infections are capable of reducing reproduction and lifespan of the most susceptible hosts, but the susceptibility to infection strongly varied between host genotypes, and most of the tested *Daphnia* clones suffered no detectable consequences of MicG infection. The host range of MicG spans across multiple *Daphnia* species; we also found it was also able to infect *Ceriodaphnia* species. Moreover, MicG was capable of infecting *D. longispina*, the host from which *O. pajunii* was described (49), which is not present in North America. Overall, our data indicates that MicG and *O. pajunii* are the same species, hence the microsporidium previously called “MicG” (4, 16, 48) should be considered *Ordospora pajunii*.

However, there are also some phenotypic differences between MicG and the European *O. pajunii*, including that spores of both *O. colligata* and *O. pajunii* isolated from Europe form chains (49), whereas MicG spores do not; this is particularly notable, as the formation of chains has been proposed as diagnostic for the genus *Ordospora* (49). In addition, European *O. pajunii* infections are much less pronounced in the hepatic caeca of the host than in the main midgut epithelium (49), and *O. colligata* frequently develops intensively in the caeca (49); in contrast, MicG infects the caeca as frequently as the midgut epithelium, and the intensity of the infection can vary temporally between the two parts of the host’s gut. Last but not least, MicG seems to have a much broader range of susceptible host species than initially found for the described isolate of *O. pajunii* and for *O. colligata* (49).

The number of host species that we observed infected (all nine of the host species studied, including those that were rare, such as *D. mendotae*) is also interesting because most microsporidia are highly host species specific (53), with only 2.2% being able to infect >4 host species (54). Prior studies on the genus *Ordospora* also suggested high specificity with regards to host species: *O. colligata* is found only in *Daphnia magna* (55), and the initial study describing *O. pajunii* found it was easily transmittable between clones of *D. longispina*, but not infectious to the two genotypes of *Daphnia magna* that were tested in lab assays (49). We confirmed MicG infections in 8 host species from two different genera in nature, and we found that it is capable of infecting allopatric *D. longispina* despite substantial geographic isolation (Table S3). The broad host range found for MicG is atypical but not unknown for microsporidia. For example, members of the genus *Encephalitozoon* are able to infect a broad spectrum of mammalian and avian hosts (56). *Encephalitozoon* spp. and *Ordospora* spp. form a common clade, indicating high genetic relatedness of the two genera; it is possible that their mechanisms of infection and interaction with host immune defense are similar. The specific determinants of infection success and host specificity in microsporidia are still being discovered, but molecular mechanisms (specifically genetic matching) seem to be the most obvious target of further investigation (53).

Broadly speaking, studies of molecular patterns of infection have provided substantial evidence for the importance of genetic compatibility between host and parasite (57). More specifically, gene-by-gene interactions have been found to determine successful infection of *Daphnia* by its parasites. For example, infections by two microsporidia, *O. colligata* and *Hamiltosporidium tvaerminnensis*, were found to be largely determined by very few host resistance genes, with some of the genes being common for infections with both parasites (58, 59). Similarly, infection of a bacterial parasite of *Daphnia*, *Pasteuria ramosa*, is determined by several genes regulating the host’s resistance (60, 61) and a few genomic regions determining the parasite’s infectivity (58, 62, 63). *P. ramosa*, similarly to MicG, is found to infect multiple *Daphnia* species (64, 65). The likelihood of multiple host species being susceptible to the same parasite species increases with increasing relatedness of the hosts (66, 67). As pathogens tend to consistently interact with the same genes across the host species (68), it is likely that high similarity of immunity genes/proteins among related host species enables a pathogen to infect multiple species. In the case of *Daphnia* susceptibility to *P. ramosa*, not only the genes but also the alleles that determine whether a particular host individual is susceptible to a given parasite strain are maintained across the *Daphnia* genus (69, 70). Hence the bacterium is infectious to multiple *Daphnia* species, including a single strain being capable of infections across host species, albeit rarely (69, 70). Given the substantial variation in susceptibility to MicG among the tested *D. dentifera* (from 0 to 100% infection prevalence) and *D. longispina* (0 to 90%) clones, and the broad range of host species of this microsporidium, it seems plausible that alleles involved in susceptibility to MicG could be conserved across host species, as in the case of *P. ramosa*. Future studies that explore the degree of host-parasite genotype specificity (as done, for example, in Carius et al. (71)), as well as the structuring of the genetic variation of the parasite by host species and geography (as done by Shaw et al. (72)) would help uncover the factors determining the likelihood of infection within and across host species.

Genetic matching between host and parasite is just one of many factors determining infection success in microsporidia (53); factors such as developmental stage and feeding behavior of the host also seem relevant to the *Daphnia* - MicG interaction. For example, some microsporidia are more infectious to their host in its early larval stages (73) whereas others are more successful infecting older life stages (74). Our results indicate that MicG is generally more infectious to younger *Daphnia*, although this effect seems to be clone-specific. The sister-microsporidian *O. colligata* was also found to be more successful infecting younger *D. magna* than older ones (75). A similar pattern of higher susceptibility of a younger host was found in *D. magna* exposed to the bacterium *P. ramosa*, and the proposed explanation was that immune defense mechanisms are less efficient in younger instars of *Daphnia* (76). In contrast, older *D. dentifera* are more susceptible to the fungus *Metschnikowia bicuspidata*; this is partially due to the higher feeding rates of older (and, therefore, larger) hosts (77), but also due to other factors, with the per spore infectivity of *M. bicuspidata* increasing as hosts age (44). Because *Daphnia* encounter parasites while foraging for phytoplankton, feeding rates determine the contact rate of the host with many parasites, including bacteria, fungi and microsporidia (75, 77–79). Even in cases where younger hosts are more susceptible to parasites, variation in feeding rates might still play an important role in determining patterns of infection. As higher contact rate with infectious spores can increase the likelihood of becoming infected (77, 78, 80), host genotypes with higher feeding rates (78, 81) can be more susceptible to parasitic infections than the slow feeders (78). Hence, variation among genotypes in feeding behavior could be another factor driving the variation in susceptibility to MicG observed among our *D. dentifera* clones.

While the infectivity of MicG largely varied between tested clones, the virulence was fairly uniform (Figure 4). We found a significant reduction in lifespan and reproduction of the host in only one genotype; the other *Daphnia* genotypes seemed largely unaffected by MicG. Low virulence could be one of the critical traits enabling MicG to generate a net positive outcome to its host when a more virulent parasite is in the environment (as found by Rogalski et al. (4)) — low basal virulence would require much less positive impact to tip the scale from parasitism to mutualism. However, two of our studies on this system found strong negative effects of MicG infection. One of them reveals a novel phenomenon of transgenerational virulence, where the offspring of hosts exposed to MicG suffered reduced fitness; this indicates that the virulence of MicG might primarily manifest in the progeny of exposed hosts (82). Another study shows that sequential exposure to MicG and then to the fungus *M. bicuspidata* reduces the lifespan and reproduction of the host, but also (due to shorter host lifespan) drastically reduces the fitness of *M. bicuspidata* (83). The latter is consistent with studies of other *Daphnia* exposed to microsporidians and then a second parasite (84, 85). Therefore, the findings of low within-generation virulence need to be interpreted cautiously.

The clone (‘S’) that was the most susceptible to MicG and suffered most from virulent effects of infection in this study also expressed the strongest transgenerational virulence effect in a related study (82). The particularly strong effect on that host may be due to two possible non-exclusive mechanisms. One possibility is that the S clone has high metabolic demands, related to its high reproduction (Figure 4C), which it has demonstrated since it was first isolated from the field. MicG damages the gut and hence likely interferes with resource assimilation, which might be harder for a clone with high resource needs to tolerate. Another reason for higher virulence in S clone could be unintended selection imposed as a result of our culturing – we maintain our lab *O. pajunii* culture in the S clone; therefore, the parasite isolate that we used in this study may have adapted to exploit this clone. While the high virulence in this host genotype might reflect selection in the lab, that does not mean that such virulence is not also occurring in nature — it is likely that each outbreak is composed of multiple MicG strains varying in their infectivity and virulence to different clones, potentially leading to selection favoring the parasite strains best adapted to exploit the most abundant hosts (62, 86). Future studies that explore variation in virulence of different *O. pajunii* isolates, as well as the potential for rapid adaptation on the part of the parasite, would help us better understand the ecological and evolutionary dynamics of this system. Moreover, it will help determine whether some of the variation in fitness impacts of infection observed in the field (4) result from variation in the virulence of the parasite due to host and/or parasite genetic variation.

The virulence of MicG might be a result of its direct impact to the host’s gut. The microsporidium frequently covers most of the midgut area in heavily infected hosts. Other studies found that *O. pajunii* reduces the penetrability of the guts of infected hosts to other parasites (4, 83), and this study found changes in the morphology of the gut (Figure 3). Most notably, microvilli were not visible next to epithelial cells with more developed infections, when the cluster had moved towards or into the gut lumen. These changes are intriguing, as they suggest that *O. pajunii* infections should strongly alter host interactions with other parasites and resources — both the reduced penetrability and the loss of microvilli should impair resource assimilation. This would be further accentuated if *Daphnia* infected with *O. pajunii* lower their feeding rate; illness mediated anorexia is common in hosts (87), including *Daphnia* infected with other parasites (78, 88). Diminished feeding rate would reduce the contact rate of the host with other parasites (thus reducing the likelihood of infection (77–79, 89)) and host nutrition (which decreases host fitness (78, 79)), making the beneficial impact of MicG contingent upon the presence of other parasites and food resources. In contrast to this prediction, in a complementary study, we found that MicG infections increase host mortality after exposure to a second pathogen that invades the host through the gut (83), which could be a consequence of overwhelming damage done to the host’s gut. Together, these impacts on gut morphology – as well as potential impacts on feeding rate – have the potential to impact not only host fitness, but also ecosystem-level processes such as primary productivity and nutrient cycling. Given that *Daphnia* are the dominant grazers in many lakes, that microsporidian transmission stages are very abundant in lakes (90), and the very high prevalence of infection in some populations (Figure 5), we propose that *O. pajunii* outbreaks might have important ecosystem-level consequences.

Our genetic analyses of the microsporidian symbiont of *Daphnia* formerly referred to as “MicG” (4, 16, 48) revealed that it belongs to *Ordospora pajunii* (49). It is found in North American lakes as well as coastal ponds in Finland, which suggests a very broad geographical distribution. It also has an intriguingly broad host distribution, infecting a wide range of *Daphnia* species as well as *Ceriodaphnia*. Our study found *O. pajunii* has modest effects on survival and reproduction within a generation, and prior work suggested context dependence of the fitness impact on hosts (4); moreover, other evidence suggests virulent effects might primarily occur in the next generation (82). Some of the virulence might relate to the destructive effects of the microsporidian on host gut morphology, with the potential for this to substantially alter interactions of this dominant herbivore with its resources. This is important because infection prevalence can become quite high, as shown in this study and in another one that matched the dynamics of *O. pajunii* spores in the water column with infections in hosts (90). Altogether, this *Daphnia*-microsporidian system seems highly promising for future studies aimed at uncovering the ecology and evolution of host-parasite interactions in the wild.

## Author contributions

- Conceptualization: MKD, KMM, ESD, TYJ, MAD
- Data curation: MKD, KMM, KS, EB, AS, SA, IVG, TYJ, MAD
- Formal analysis: MKD, KMM, KS, EB, AW, SA, IVG, TYJ, MAD
- Funding acquisition: IVH, TYJ, MAD
- Investigation: MKD, KMM, KS, ESD, MAR, CDG, EB, MV, CH, RJ, FEC, AW, SA, AS, MN, ST, KB, IVG, TYJ, MAD
- Methodology: MKD, KMM, KS, ESD, AW, SA, AS, MN, ST, KB, IVG, TYJ, MAD
- Project administration: KB, IVG, TYJ, MAD
- Resources: AW, IVG, TYJ, MAD
- Software:
- Supervision: MKD, KMM, IVG, TYJ, MAD
- Validation: MKD, KMM, KS, AW, SA, AS, MN, ST, KB, IVG, TYJ, MAD
- Visualization: MKD, KMM, KS, AW, TYJ, MAD
- Writing - original draft: MKD, KMM, TYJ, MAD
- Writing - review & editing: MKD, KMM, KS, ESD, MAR, CDG, EB, MV, CH, RJ, FEC, AW, SA, AS, MN, ST, KB, IVG, TYJ, MAD

## Acknowledgements

We thank Justyna Wolinska for the *D. longispina* clones and Dylan Baker for the Lake Erie sample. We also thank the members of the Duffy Lab who helped collect and process the field samples during the host range portion of this study – Rebecca Bilich, Siobhan Calhoun, Kit McLean, Kira Monell, Kristel Sánchez, Teresa Sauer, and Syuan-Jyun Sun. We owe thanks to the anonymous reviewers, whose input helped us improve the manuscript. This work was supported by NSF grants DEB-1305836 and DEB-1748729 and by the Gordon and Betty Moore Foundation (GBMF9202; DOI: https://doi.org/10.37807/GBMF9202). The work (proposal 10.46936/10.25585/60001004) conducted by the U.S. Department of Energy Joint Genome Institute (https://ror.org/04xm1d337), a DOE Office of Science User Facility, is supported by the Office of Science of the U.S. Department of Energy under Contract No. DE-AC02-05CH11231. This project was funded in part by the National Science Foundation grant DEB-1929738 to TYJ. TYJ is a fellow of the Canadian Institute for Advanced Research program Fungal Kingdom: Threats & Opportunities. KS was supported by JSPS Overseas Research Fellowships (No. 201960485).

## References

1. Keeling Patrick J., Slamovits Claudio H. 2004. Simplicity and complexity of microsporidian genomes. Eukaryot Cell 3:1363–1369.

2. Dean P, Hirt RP, Embley TM. 2016. Microsporidia: why make nucleotides if you can steal them? PLOS Pathog 12:e1005870.

3. Bojko J, Reinke AW, Stentiford GD, Williams B, Rogers MSJ, Bass D. 2022. Microsporidia: a new taxonomic, evolutionary, and ecological synthesis. Trends Parasitol 38:642–659.

4. Rogalski MA, Merrill TS, Gowler C, Caceres C, Duffy M. 2021. Context-dependent host-symbiont interactions: shifts along the parasitism–mutualism continuum. Am Nat 198:563–575.

5. Ryan JA, Kohler SL. 2010. Virulence is context-dependent in a vertically transmitted aquatic host–microparasite system. Int J Parasitol 40:1665–1673.

6. Vilcinskas A, Stoecker K, Schmidtberg H, Röhrich CR, Vogel H. 2013. Invasive harlequin ladybird carries biological weapons against native competitors. Science 340:862–863.

7. Shi W, Guo Y, Xu C, Tan S, Miao J, Feng Y, Zhao H, St. Leger RJ, Fang W. 2014. Unveiling the mechanism by which microsporidian parasites prevent locust swarm behavior. Proc Natl Acad Sci 111:1343–1348.

8. Herren JK, Mbaisi L, Mararo E, Makhulu EE, Mobegi VA, Butungi H, Mancini MV, Oundo JW, Teal ET, Pinaud S, Lawniczak MKN, Jabara J, Nattoh G, Sinkins SP. 2020. A microsporidian impairs *Plasmodium falciparum* transmission in *Anopheles arabiensis* mosquitoes. Nat Commun 11:2187.

9. Fries I. 1993. *Nosema Apis*—A parasite in the honey bee colony. Bee World 74:5–19.

10. Lewis LC, Bruck DJ, Prasifka JR, Raun ES. 2009. *Nosema pyrausta*: Its biology, history, and potential role in a landscape of transgenic insecticidal crops. Biol Control 48:223–231.

11. Vávra J, Lukeš J. 2013. Chapter Four - Microsporidia and ‘The Art of Living Together,’ p. 253–319. In Rollinson, D (ed.), Advances in Parasitology. Academic Press.

12. Stentiford GD, Becnel J.J., Weiss LM, Keeling PJ, Didier ES, Williams BAP, Bjornson S, Kent ML, Freeman MA, Brown MJF, Troemel ER, Roesel K, Sokolova Y, Snowden KF, Solter L. 2016. Microsporidia – emergent pathogens in the global food chain. Trends Parasitol 32:336–348.

13. Ewald PW. 1987. Transmission modes and evolution of the parasitism-mutualism continuuma. Ann N Y Acad Sci 503:295–306.

14. Lindsay RJ, Holder PJ, Talbot NJ, Gudelj I. 2023. Metabolic efficiency reshapes the seminal relationship between pathogen growth rate and virulence. Ecol Lett 26:896–907.

15. Drew GC, Stevens EJ, King KC. 2021. Microbial evolution and transitions along the parasite–mutualist continuum. Nat Rev Microbiol 19:623–638.

16. Westphal GH, Stewart Merrill TE. 2022. Partitioning variance in immune traits in a zooplankton host—Fungal parasite system. Ecol Evol 12:e9640.

17. Seda J, Petrusek A. 2011. *Daphnia* as a model organism in limnology and aquatic biology: introductory remarks. J Limnol 70:337–344.

18. Miner BE, De Meester L, Pfrender ME, Lampert W, Hairston Jr NG. 2012. Linking genes to communities and ecosystems: *Daphnia* as an ecogenomic model. Proc R Soc B 279:1873–1882

19. Anderson RM, May RM. 1981. The population dynamics of microparasites and their invertebrate hosts. Philos Trans R Soc Lond B Biol Sci 291:451–524.

20. Bedhomme S, Agnew P, Vital Y, Sidobre C, Michalakis Y. 2005. Prevalence-dependent costs of parasite virulence. PLOS Biol 3:e262.

21. Tessier AJ, Woodruff P. 2002. Cryptic trophic cascade along a gradient of lake size. Ecology 83:1263–1270.

22. Duffy MA, Caceres CE, Hall SR, Tessier AJ, Ives AR. 2010. Temporal, spatial, and between-host comparisons of patterns of parasitism in lake zooplankton. Ecology 91:3322–3331.

23. Duffy Meghan A., James Timothy Y., Longworth Alan, Drake H. L. 2015. Ecology, virulence, and phylogeny of *Blastulidium paedophthorum*, a widespread brood parasite of *Daphnia* spp. Appl Environ Microbiol 81:5486–5496.

24. Trzebny A, Slodkowicz-Kowalska A, Becnel JJ, Sanscrainte N, Dabert M. 2020. A new method of metabarcoding Microsporidia and their hosts reveals high levels of microsporidian infections in mosquitoes (Culicidae). Mol Ecol Resour 20:1486–1504.

25. Kumar S, Stecher G, Li M, Knyaz C, Tamura K. 2018. MEGA X: Molecular Evolutionary Genetics Analysis across computing platforms. Mol Biol Evol 35:1547–1549.

26. Keeling PJ. 2014. Phylogenetic place of Microsporidia in the tree of eukaryotes, pp. 195–202 In Weiss L, Becnel JJ, editors. Microsporidia: Pathogens of Opportunity. John Wiley & Sons.

27. James TY, Stajich JE, Hittinger CT, Rokas A. 2020. Toward a fully resolved fungal tree of life. Annu Rev Microbiol 74:291–313.

28. Schoch CL, Seifert KA, Huhndorf S, Robert V, Spouge JL, Levesque CA, Chen W, null null, null null, Bolchacova E, Voigt K, Crous PW, Miller AN, Wingfield MJ, Aime MC, An K-D, Bai F-Y, Barreto RW, Begerow D, Bergeron M-J, Blackwell M, Boekhout T, Bogale M, Boonyuen N, Burgaz AR, Buyck B, Cai L, Cai Q, Cardinali G, Chaverri P, Coppins BJ, Crespo A, Cubas P, Cummings C, Damm U, de Beer ZW, de Hoog GS, Del-Prado R, Dentinger B, Diéguez-Uribeondo J, Divakar PK, Douglas B, Dueñas M, Duong TA, Eberhardt U, Edwards JE, Elshahed MS, Fliegerova K, Furtado M, García MA, Ge Z-W, Griffith GW, Griffiths K, Groenewald JZ, Groenewald M, Grube M, Gryzenhout M, Guo L-D, Hagen F, Hambleton S, Hamelin RC, Hansen K, Harrold P, Heller G, Herrera C, Hirayama K, Hirooka Y, Ho H-M, Hoffmann K, Hofstetter V, Högnabba F, Hollingsworth PM, Hong S-B, Hosaka K, Houbraken J, Hughes K, Huhtinen S, Hyde KD, James T, Johnson EM, Johnson JE, Johnston PR, Jones EBG, Kelly LJ, Kirk PM, Knapp DG, Kõljalg U, Kovács GM, Kurtzman CP, Landvik S, Leavitt SD, Liggenstoffer AS, Liimatainen K, Lombard L, Luangsa-ard JJ, Lumbsch HT, Maganti H, Maharachchikumbura SSN, Martin MP, May TW, McTaggart AR, Methven AS, Meyer W, Moncalvo J-M, Mongkolsamrit S, Nagy LG, Nilsson RH, Niskanen T, Nyilasi I, Okada G, Okane I, Olariaga I, Otte J, Papp T, Park D, Petkovits T, Pino-Bodas R, Quaedvlieg W, Raja HA, Redecker D, Rintoul TL, Ruibal C, Sarmiento-Ramírez JM, Schmitt I, Schüßler A, Shearer C, Sotome K, Stefani FOP, Stenroos S, Stielow B, Stockinger H, Suetrong S, Suh S-O, Sung G-H, Suzuki M, Tanaka K, Tedersoo L, Telleria MT, Tretter E, Untereiner WA, Urbina H, Vágvölgyi C, Vialle A, Vu TD, Walther G, Wang Q-M, Wang Y, Weir BS, Weiß M, White MM, Xu J, Yahr R, Yang ZL, Yurkov A, Zamora J-C, Zhang N, Zhuang W-Y, Schindel D. 2012. Nuclear ribosomal internal transcribed spacer (ITS) region as a universal DNA barcode marker for Fungi. Proc Natl Acad Sci 109:6241–6246.

29. Davis WJ, Amses KR, Benny GL, Carter-House D, Chang Y, Grigoriev I, Smith ME, Spatafora JW, Stajich JE, James TY. 2019. Genome-scale phylogenetics reveals a monophyletic Zoopagales (Zoopagomycota, Fungi). Mol Phylogenet Evol 133:152–163.

30. Bankevich A, Nurk S, Antipov D, Gurevich AA, Dvorkin M, Kulikov AS, Lesin VM, Nikolenko SI, Pham S, Prjibelski AD, Pyshkin AV, Sirotkin AV, Vyahhi N, Tesler G, Alekseyev MA, Pevzner PA. 2012. SPAdes: A new genome assembly algorithm and its applications to single-cell sequencing. J Comput Biol 19:455–477.

31. Grigoriev IV, Nikitin R, Haridas S, Kuo A, Ohm R, Otillar R, Riley R, Salamov A, Zhao X, Korzeniewski F, Smirnova T, Nordberg H, Dubchak I, Shabalov I. 2014. MycoCosm portal: gearing up for 1000 fungal genomes. Nucleic Acids Res 42:D699–D704.

32. Manni M, Berkeley MR, Seppey M, Simão FA, Zdobnov EM. 2021. BUSCO Update: Novel and streamlined workflows along with broader and deeper phylogenetic coverage for scoring of eukaryotic, prokaryotic, and viral genomes. Mol Biol Evol 38:4647–4654.

33. McGowan J, O’Hanlon R, Owens RA, Fitzpatrick DA. 2020. Comparative genomic and proteomic analyses of three widespread *Phytophthora* species: *Phytophthora chlamydospora*, *Phytophthora gonapodyides* and *Phytophthora pseudosyringae*. Microorganisms 8:653.

34. Edgar RC. 2004. MUSCLE: multiple sequence alignment with high accuracy and high throughput. Nucleic Acids Res 32:1792–1797.

35. Capella-Gutiérrez S, Silla-Martínez JM, Gabaldón T. 2009. trimAl: a tool for automated alignment trimming in large-scale phylogenetic analyses. Bioinformatics 25:1972–1973.

36. Minh BQ, Schmidt HA, Chernomor O, Schrempf D, Woodhams MD, von Haeseler A, Lanfear R. 2020. IQ-TREE 2: New models and efficient methods for phylogenetic inference in the genomic era. Mol Biol Evol 37:1530–1534.

37. Wang H-C, Minh BQ, Susko E, Roger AJ. 2018. Modeling site heterogeneity with posterior mean site frequency profiles accelerates accurate phylogenomic estimation. Syst Biol 67:216–235.

38. Chernomor O, von Haeseler A, Minh BQ. 2016. Terrace aware data structure for phylogenomic inference from supermatrices. Syst Biol 65:997–1008.

39. Lee I, Ouk Kim Y, Park S-C, Chun J. 2016. OrthoANI: An improved algorithm and software for calculating average nucleotide identity. Int J Syst Evol Microbiol 66:1100–1103.

40. Treangen TJ, Ondov BD, Koren S, Phillippy AM. 2014. The Harvest suite for rapid core-genome alignment and visualization of thousands of intraspecific microbial genomes. Genome Biol 15:524.

41. Cingolani P, Platts A, Wang LL, Coon M, Nguyen T, Wang L, Land SJ, Lu X, Ruden DM. 2012. A program for annotating and predicting the effects of single nucleotide polymorphisms, SnpEff. Fly (Austin) 6:80–92.

42. Hyatt D, Chen G-L, LoCascio PF, Land ML, Larimer FW, Hauser LJ. 2010. Prodigal: prokaryotic gene recognition and translation initiation site identification. BMC Bioinformatics 11:119.

43. Emms DM, Kelly S. 2019. OrthoFinder: phylogenetic orthology inference for comparative genomics. Genome Biol 20:238.

44. Clay PA, Gattis S, Garcia J, Hernandez V, Ben-Ami F, Duffy MA. 2023. Age structure eliminates the impact of coinfection on epidemic dynamics in a freshwater zooplankton system. Am Nat 202:785–799.

45. Ben-Ami F. 2019. Host age effects in invertebrates: epidemiological, ecological, and evolutionary implications. Trends Parasitol 35:466–480.

46. Duffy MA, Hall SR. 2008. Selective predation and rapid evolution can jointly dampen effects of virulent parasites on *Daphnia* populations. Am Nat 171:499–510.

47. Shaw CL, Hall SR, Overholt EP, Cáceres CE, Williamson CE, Duffy MA. 2020. Shedding light on environmentally transmitted parasites: lighter conditions within lakes restrict epidemic size. Ecology 101:e03168.

48. Gowler CD, Rogalski MA, Shaw CL, Hunsberger KK, Duffy MA. 2021. Density, parasitism, and sexual reproduction are strongly correlated in lake *Daphnia* populations. Ecol Evol 11:10446–10456.

49. de Albuquerque NRM, Haag KL, Fields PD, Cabalzar A, Ben-Ami F, Pombert J-F, Ebert D. 2022. A new microsporidian parasite, Ordospora pajunii sp. nov (Ordosporidae), of Daphnia longispina highlights the value of genomic data for delineating species boundaries. J Eukaryot Microbiol 69:e12902.

50. Dia N, Lavie L, Faye N, Méténier G, Yeramian E, Duroure C, Toguebaye BS, Frutos R, Niang MN, Vivarès CP, Ben Mamoun C, Cornillot E. 2016. Subtelomere organization in the genome of the microsporidian *Encephalitozoon cuniculi*: patterns of repeated sequences and physicochemical signatures. BMC Genomics 17:34.

51. Krzywinski M, Schein J, Birol İ, Connors J, Gascoyne R, Horsman D, Jones SJ, Marra MA. 2009. Circos: An information aesthetic for comparative genomics. Genome Res 19:1639– 1645.

52. Ebert D. 2022. *Daphnia* as a versatile model system in ecology and evolution. EvoDevo 13:16.

53. Willis AR, Reinke AW. 2022. Factors that determine microsporidia infection and host specificity, p. 91–114. In Weiss, LM, Reinke, AW (eds.), Microsporidia: Current Advances in Biology. Springer International Publishing, Cham.

54. Murareanu Brandon M., Sukhdeo Ronesh, Qu Rui, Jiang Jason, Reinke Aaron W. 2021. Generation of a Microsporidia species attribute database and analysis of the extensive ecological and phenotypic diversity of Microsporidia. mBio 12:e01490–21.

55. Larsson JIR, Ebert D, Vávra J. 1997. Ultrastructural study and description of *Ordospora colligata* gen. et sp. nov. (microspora, ordosporidae fam. nov.), a new microsporidian parasite of *Daphnia magna* (Crustacea, Cladocera). Eur J Protistol 33:432–443.

56. Hinney B, Sak B, Joachim A, Kváč M. 2016. More than a rabbit’s tale – *Encephalitozoon* spp. in wild mammals and birds. Artic 25th WAAVP Conf Liverp August 2015 5:76–87.

57. Dybdahl MF, Jenkins CE, Nuismer SL. 2014. Identifying the molecular basis of host-parasite coevolution: merging models and mechanisms. Am Nat 184:1–13.

58. Routtu J, Ebert D. 2015. Genetic architecture of resistance in *Daphnia* hosts against two species of host-specific parasites. Heredity 114:241–248.

59. Keller D, Kirk D, Luijckx P. 2019. Four QTL underlie resistance to a microsporidian parasite that may drive genome evolution in its *Daphnia* host. bioRxiv 847194.

60. Ameline C, Voegtli F, Andras J, Dexter E, Engelstädter J, Ebert D. Genetic slippage after sex maintains diversity for parasite resistance in a natural host population. Sci Adv 8:eabn0051.

61. Ameline C, Bourgeois Y, Vögtli F, Savola E, Andras J, Engelstädter J, Ebert D. 2021. A two-locus system with strong epistasis underlies rapid parasite-mediated evolution of host resistance. Mol Biol Evol 38:1512–1528.

62. Andras JP, Fields PD, Du Pasquier L, Fredericksen M, Ebert D. 2020. genome-wide association analysis identifies a genetic basis of infectivity in a model bacterial pathogen. Mol Biol Evol 37:3439–3452.

63. Alix Thivolle, Marjut Paljakka, Dieter Ebert, Peter D. Fields. 2024. The genome of *Pasteuria ramosa* reveals a high turnover rate of collagen-like genes. bioRxiv 2024.02.09.579640.

64. Ebert D, Rainey P, Embley TM, Scholz D. 1996. Development, life cycle, ultrastructure and phylogenetic position of *Pasteuria ramosa* Metchnikoff 1888: rediscovery of an obligate endoparasite of *Daphnia magna* Straus. Philos Trans R Soc Lond B Biol Sci 351:1689– 1701.

65. Shaw CL. 2019. Drivers of epidemic timing and size in a natural aquatic system. Doctoral dissertation. University of Michigan, Michigan, USA. https://deepblue.lib.umich.edu/handle/2027.42/151685.

66. Davies TJ, Pedersen AB. 2008. Phylogeny and geography predict pathogen community similarity in wild primates and humans. Proc R Soc B Biol Sci 275:1695–1701.

67. Parker IM, Saunders M, Bontrager M, Weitz AP, Hendricks R, Magarey R, Suiter K, Gilbert GS. 2015. Phylogenetic structure and host abundance drive disease pressure in communities. Nature 520:542–544.

68. Shultz AJ, Sackton TB. 2019. Immune genes are hotspots of shared positive selection across birds and mammals. eLife 8:e41815.

69. Luijckx P, Duneau D, Andras JP, Ebert D. 2014. Cross-species infection trials reveal cryptic parasite varieties and a putative polymorphism shared among host species. Evolution 68:577–586.

70. Duneau D, Luijckx P, Ben-Ami F, Laforsch C, Ebert D. 2011. Resolving the infection process reveals striking differences in the contribution of environment, genetics and phylogeny to host-parasite interactions. BMC Biol 9:11.

71. Carius HJ, Little TJ, Ebert D. 2001. Genetic variation in a host-parasite association: potential for coevolution and frequency-dependent selection. Evolution 1136–1145.

72. Shaw CL, Bilich R, O’Brien B, Cáceres CE, Hall SR, James TY, Duffy MA. 2021. Genotypic variation in an ecologically important parasite is associated with host species, lake and spore size. Parasitology, 2021/06/09 ed. 148:1303–1312.

73. Blaser M, Schmid-Hempel P. 2005. Determinants of virulence for the parasite *Nosema whitei* in its host *Tribolium castaneum*. J Invertebr Pathol 89:251–257.

74. Balla KM, Andersen EC, Kruglyak L, Troemel ER. 2015. A wild *C. elegans* strain has enhanced epithelial immunity to a natural microsporidian parasite. PLOS Pathog 11:e1004583.

75. Kirk D, Luijckx P, Stanic A, Krkošek M. 2019. predicting the thermal and allometric dependencies of disease transmission via the metabolic theory of ecology. Am Nat 193:661–676.

76. Izhar R, Ben-Ami F. 2015. Host age modulates parasite infectivity, virulence and reproduction. J Anim Ecol 84:1018–1028.

77. Hall SR, Sivars-Becker L, Becker C, Duffy MA, Tessier AJ, Cáceres CE. 2007. Eating yourself sick: transmission of disease as a function of foraging ecology. Ecol Lett 10:207– 218.

78. Auld SKJR, Raidma K. 2022. Host genetic variation in feeding rate mediates a fecundity cost of parasite resistance in a *Daphnia*-parasite system. bioRxiv 2022.11.29.518345.

79. Hall SR, Becker CR, Duffy MA, Cáceres CE. 2010. Variation in resource acquisition and use among host clones creates key epidemiological trade-offs. Am Nat 176:557–565.

80. Shocket MS, Vergara D, Sickbert AJ, Walsman JM, Strauss AT, Hite JL, Duffy MA, Cáceres CE, Hall SR. 2018. Parasite rearing and infection temperatures jointly influence disease transmission and shape seasonality of epidemics. Ecology 99:1975–1987.

81. Hite JL, Pfenning-Butterworth AC, Vetter RE, Cressler CE. 2020. A high-throughput method to quantify feeding rates in aquatic organisms: A case study with *Daphnia*. Ecol Evol 10:6239–6245.

82. McIntire KM, Dziuba MK, Haywood EB, Robertson ML, Vaandrager M, Baird E, Corcoran F, Cortez MH, Duffy MA. 2023. Transgenerational virulence: Maternal pathogen exposure reduces offspring fitness. bioRxiv 2023.03.14.532659.

83. Marcin K. Dziuba, Kristina M. McIntire, Elizabeth S. Davenport, Emma Baird, Cristian Huerta, Riley Jaye, Fiona Corcoran, Paige McCreadie, Taleah Nelson, Meghan A. Duffy. 2024. Microsporidian coinfection reduces fitness of a fungal pathogen due to rapid host mortality. bioRxiv 2024.02.08.579564.

84. Manzi F, Halle S, Seemann L, Ben-Ami F, Wolinska J. 2021. Sequential infection of *Daphnia magna* by a gut microsporidium followed by a haemolymph yeast decreases transmission of both parasites. Parasitology 1–42.

85. Holland CV, Luijckx P, O’Keeffe FE, Pendleton RC. 2023. Increased virulence due to multiple infection in *Daphnia* leads to limited growth in 1 of 2 co-infecting microsporidian parasites. Parasitology 151:58–67.

86. Lively CM, Dybdahl MF. 2000. Parasite adaptation to locally common host genotypes. Nature 405:679–681.

87. Hite JL, Pfenning AC, Cressler CE. 2020. Starving the enemy? Feeding behavior shapes host-parasite interactions. Trends Ecol Evol 35:68–80.

88. Penczykowski RM, Shocket MS, Ochs JH, Lemanski BCP, Sundar H, Duffy MA, Hall SR. 2022. Virulent disease epidemics can increase host density by depressing foraging of hosts. Am Nat 199:75–90.

89. Shocket MS, Strauss AT, Hite JL, Šljivar M, Civitello DJ, Duffy MA, Cáceres CE, Hall SR. 2018. Temperature drives epidemics in a zooplankton-fungus disease system: A trait-driven approach points to transmission via host foraging. Am Nat 191:435–451.

90. Davenport ES, Dziuba MK, Jacobson LE, Calhoun SK, Monell KJ, Duffy MA. 2024. How does parasite environmental transmission stage concentration change before, during, and after disease outbreaks? Ecology 105:e4235.

