## Supplemental Materials for "Phylogeny, morphology, virulence, ecology, and host range of *Ordospora pajunii* (Ordosporidae), a microsporidian symbiont of *Daphnia* spp"

#### Cultivating MicG in the laboratory

*Establishing lab cultures*

We established protocols for maintaining MicG in the *D. dentifera* clone S. These cultures were established in water from North Lake (Washtenaw County, Michigan, USA) that was filtered (with Pall AE glass microfiber filters) in order to remove algae, cyanobacteria, and debris from the water. To establish the MicG culture, we collected *D. dentifera* visibly infected with MicG from Walsh Lake (Washtenaw County, Michigan, USA). We grouped >10 individuals in a 1.5mL centrifuge tube and ground them using a plastic pestle. We filled 24 centrifuge tubes (1.5mL volume capacity), each with 1mL of filtered lake water and introduced a single 0-2 day old *D. dentifera* (S clone) to each tube. Individuals were left in their tubes for 48h to ingest the parasite spores. After two days we transferred each host individual to a separate 50mL beaker, which we fed 20,000 cells/mL of *Ankistrodesmus falcatus* 3 times a week. We screened for infection after seven days, then checked at 3 day intervals. Each individual that was found to be infected was used to establish a separate 'farm'. New farms were started by moving a heavily infected *Daphnia* (i.e. *Daphnia* having tens to hundreds of microsporidian spore clusters) into a 300mL beaker and adding 5 juveniles of the S clone. As the MicG farm grew, we increased its volume to 1L.

*Culture observations*

Initially the number of infected *Daphnia* increases as the host population size increases. Due to the near-constant shedding of spores, host offspring quickly become infected, increasing the number of infected hosts. Soon after the host population size approaches the carrying capacity of the farm, the infection prevalence among adults frequently reaches 90-100%. In high prevalence farms, we tend to observe infection symptoms even in juveniles. We hypothesize that this is due to high spore concentrations in the environment. If prevalence is low, infections are infrequent in juveniles. The demography of the farm quickly shifts as the adults start producing non-viable embryos or stop producing embryos. Shortly thereafter, the host population collapses.

#### Molecular analyses

*Accession numbers*

**SSU: OQ754180**
**ITS: OQ754181**
**Whole genome: SRR24010758**

### A) Alpha/beta hydrolase (KAH9410807.1)

```
Consensus                  MEXEFENKLNSTNIIDKXSGIIMIICALSIWIAIFGVCLLLIVVVIFKQFEYYIIIFQTR   60
MicG_NODE_16_21_hydrolase ..S..............V..........................................   60
Op_KAH9410807.1_hydrolase ..A..............A..........................................   60

Consensus                  AVNLXXSFLDLXKIYPXKSQXLLLFEILIGVYKCNPHYDDKNQTFTYDIYDNKSNKITNY  120
MicG_NODE_16_21_hydrolase ....FC.....K....E...E.......................................  120
Op_KAH9410807.1_hydrolase ....LY.....T....K...S.......................................  120

Consensus                  IPMEDFSVISGQGSDICMLGXNHLKENGKLIVFFPGNAYHLIAVLEIIKXYFERYKSDKV  180
MicG_NODE_16_21_hydrolase ....................K............................E..........  180
Op_KAH9410807.1_hydrolase ....................E............................K..........  180

Consensus                  CVAAMIYRGMNESLTPSEKNLKNDIQNFKRIIKETNPNTKFIIYGFSIGCAPAIHLASII  240

Consensus                  IYLFISEXDBXIKPKTNQLVLXNLLSEKRTLFINFMPKIVKTPRXXSRDXSXHNIDMSTD  360
MicG_NODE_16_21_hydrolase .......Y.DL..........S......................NH...Y.P........  360
Op_KAH9410807.1_hydrolase .......D.NV..........N......................TL...C.R........  360

Consensus                  INEIINASIRHNK                                                 373
MicG_NODE_16_21_hydrolase .............                                                 373
Op_KAH9410807.1_hydrolase .............                                                 373
```

### B) Hypothetical protein (KAH9411254.1)

```
Consensus           MTSPITVGKSPITVRNLELDDQTKIEEALSIGMAKKFVNHTFKRVDGISSLIFFVLGGLY   60
MicG_NODE_18_24_hyp ............................................................   60
Op_KAH9411254.1_hyp ............................................................   60

Consensus           TLISVFTEFMSEGRLLEVLNTDNVVKAFICIPIVLMLLKIIYLMKKHLSAFGFKQVSTGK  120
MicG_NODE_18_24_hyp ............................................................  120
Op_KAH9411254.1_hyp ............................................................  120

Consensus           FIVMSLYLLTLLLMLLLIISGKLKFDRKAVPIGISIMAFFSMVVSNEFVEFRKQKDRNYV  180
MicG_NODE_18_24_hyp ............................................................  180
Op_KAH9411254.1_hyp ............................................................  180

Consensus           DYFIIIGAAFITIDWILRIVSVVLDAYYEXRAITXXNAAWYLKLESYSNGFHMTTIFLLL  240
MicG_NODE_18_24_hyp .............................E....PD........................  240
Op_KAH9411254.1_hyp .............................K....SE........................  240

Consensus           AFPLIQNFLCEVDALFEETTDEQFQILNVLCIIALVFGILAIIMKAEIFSKHISXKISXK  300
MicG_NODE_18_24_hyp ......................................................E...A.  300
Op_KAH9411254.1_hyp ......................................................K...D.  300

Consensus           FIXRLXTLPSXXX                                                 313
MicG_NODE_18_24_hyp ..D..K....GGN                                                 313
Op_KAH9411254.1_hyp ..A..N....YDK                                                 313
```

### C) Helicase (KAH9411242.1)

```
Consensus                MYISYAEYVNMIANMVGEXDEGKLERMVGALLSCDASGEQLCRKVEEAIGKRIAPDVVDG    60
MicG_NODE_18_11_helicase ..................S.........................................    60
Op_KAH9411242.1_helicase ..................G.........................................    60

Consensus                LARLTQSFCGNDRSSVLFEDVELFDALFASDEKRMLLEVIRGGVLDRLCVLLRGWKMYFL   120
MicG_NODE_18_11_helicase ............................................................   120
Op_KAH9411242.1_helicase ............................................................   120

Consensus                NRILCNSKAFEYFLVYQLEPEDSMKAKEMLRTEGAVAFTKYIEHDMLSLMSGCKIVDLER   180
MicG_NODE_18_11_helicase ............................................................   180
Op_KAH9411242.1_helicase ............................................................   180

Consensus                TSSNASTHMKKREYPADAEVRYADGIETVYVHGCKKDIDFDADVPGDAELLFGNGFKFNY   240
MicG_NODE_18_11_helicase ............................................................   240
Op_KAH9411242.1_helicase ............................................................   240

Consensus                VQSAVYDAVFNDSGNVLVCAPTGSGKTVIGALSIXKVVAQQGKMGDAKNAGADGGKKIAY   300
MicG_NODE_18_11_helicase ..................................F.........................   300
Op_KAH9411242.1_helicase ..................................L.........................   300

Consensus                VVPMRALAREVSMTLGQMFSRHGMAVVEHTSDTDVGYKHLEKADVIVCTPEKLDALTRNT   360
MicG_NODE_18_11_helicase ............................................................   360
Op_KAH9411242.1_helicase ............................................................   360

Consensus                GFGFSVMVIDEIHMLDEDRGATIEALVARMSVRKGCRIIGLSATLPNHMDVGRFLGCKQE   420
MicG_NODE_18_11_helicase ............................................................   420
Op_KAH9411242.1_helicase ............................................................   420

Consensus                SVFAFGSEFRRCAMDYELINVGMREMEMIVAVEKVLENMDIGGPVIVFVHSRRDVMEMAR   480
MicG_NODE_18_11_helicase ............................................................   480
Op_KAH9411242.1_helicase ............................................................   480

Consensus                ELARYLGRHECRDNDDDKLLADAEKSIREIARHGIGVHHAGLSRKVRTAMEELYKSGRID   540
MicG_NODE_18_11_helicase ............................................................   540
Op_KAH9411242.1_helicase ............................................................   540

Consensus                VMVCTATLAWGVNLPGRTVVIKGADVYDTYTCEWKSIKQAEILQMFGRAGRFGDERCKGV   600
MicG_NODE_18_11_helicase ............................................................   600
Op_KAH9411242.1_helicase ............................................................   600

Consensus                LVSSKQTEFLVERCIESRLLPRLCDFLNAEIAGGMKEFGQAVDWFKHTFYYARLVVMNRE   660
MicG_NODE_18_11_helicase ............................................................   660
Op_KAH9411242.1_helicase ............................................................   660

Consensus                SGKAAREIVYSALKHLEAAGLIVFEPFVHSTCVGAVASRYCLXYRDSSRMFADLCSLMTD   720
MicG_NODE_18_11_helicase ..........................................H.................   720
Op_KAH9411242.1_helicase ..........................................Q.................   720

Consensus                SSMFSMIAEMSEFKGIGIXKEEFEAGDAVNMVPVPTESTFGILVQCHVANRVELTVLSQN   780
MicG_NODE_18_11_helicase ..................L.........................................   780
Op_KAH9411242.1_helicase ..................R.........................................   780

Consensus                LPRMFAALFDVSVRKGLGVCRRVIRWYKAAVHRIFPYQTPLRHFCTNHDALRMLEMKEIP   840
```

```
Consensus                SDDRYYVLITDSRDKELAVCDTVVFGDGMHSSQNYAINTKSPFLNTYVLSANYLCPEDPS   960
MicG_NODE_18_11_helicase ............................................................   960
Op_KAH9411242.1_helicase ............................................................   960

Consensus                VLSLVEAGNAERSMFSSVWREFMIDVLSRSRGNVVRLGLVDGAICTKTVIVAGYRERRRL  1020
MicG_NODE_18_11_helicase ............................................................  1020
Op_KAH9411242.1_helicase ............................................................  1020

Consensus                RMMGYEAYVHDEFVSGRVTCESVTIMDVHNIFSNYLIEACIAQCIVQDIRMVLVGLPFVD  1080
MicG_NODE_18_11_helicase ............................................................  1080
Op_KAH9411242.1_helicase ............................................................  1080

Consensus                GADIEHLGDIEQPGEADRSKSGCNECKEKHHMEYEMKEKTRISMYGSCSLTHHGEMHDYL  1140
MicG_NODE_18_11_helicase ............................................................  1140
Op_KAH9411242.1_helicase ............................................................  1140

Consensus                NEFELNXSRMSVDGSCLVILPVQSACRHFASRFRDARIAAEFGCIGKGVHFATRRSIDMW  1200
MicG_NODE_18_11_helicase ......M.....................................................  1200
Op_KAH9411242.1_helicase ......I.....................................................  1200

Consensus                IDAGWNPCVEQVHVIGTVYYDHESQMYADYRVADVCRYSMLGSRVFIYTKRSKALLYLNQ  1260
MicG_NODE_18_11_helicase ............................................................  1260
Op_KAH9411242.1_helicase ............................................................  1260

Consensus                GRIPLHYRSSGRGELDVYSYWIGCNLNKXVACTSLLVDENGLTRYGRILCRYGMCIDTIR  1320
MicG_NODE_18_11_helicase ............................R...............................  1320
Op_KAH9411242.1_helicase ............................H...............................  1320

Consensus                MFVGSVKDKMGLKNILLLVCRAEELVFCMDHDEYKVLSSNGVDISRGKAYGLASHDFVNE  1380
MicG_NODE_18_11_helicase ............................................................  1380
Op_KAH9411242.1_helicase ............................................................  1380

Consensus                SECPDILQYYLDDVLPIIYKMYLCLLEVSIEKACLKTAFNVMFGLQSITKKTCAAQDKFY  1440
MicG_NODE_18_11_helicase ............................................................  1440
Op_KAH9411242.1_helicase ............................................................  1440

Consensus                KLKLVENSVVIDVASMPDAPVGFVVFFLFSGSNEYEILKVESPGXYKVEWRSRACYVVSD  1500
MicG_NODE_18_11_helicase ............................................V...............  1500
Op_KAH9411242.1_helicase ............................................M...............  1500

Consensus                CCTGFDAIGCELTDGNASV                                           1519
MicG_NODE_18_11_helicase ...................                                           1519
Op_KAH9411242.1_helicase ...................                                           1519
```

### D) Aminopeptidase (KAH9410533.1), note MicG protein is partial

```
Consensus                         MXANHVIAAFLLYMCSHTQSIHRSHRYSIYTKYRCMKGMYGYAMASQAQSIHNAQHIQHT    60
MicG_NODE_26_1_peptidase_partial  .G..........................................................    60
Op_KAH9410533.1_peptidase         .R..........................................................    60

Consensus                         HHKYIYSNDGTNSSHTGSIGTDDVLHANDVLHINXREVLGSNVVPQHYDLRIKIMDAGFS   120
MicG_NODE_26_1_peptidase_partial  ..................................D.........................   120
Op_KAH9410533.1_peptidase         ..................................Y.........................   120

Consensus                         GAVGIAIDIKNATRSIVLNAAGLDVSVAKIVAGGRHHTGVVHSGMQLDVRDQIRIEFAEH   180
MicG_NODE_26_1_peptidase_partial  ............................................................   180
Op_KAH9410533.1_peptidase         ............................................................   180

Consensus                         VKGPAYILLQFSGAYATGLRGLYXSNACTSXXNGNDNLNGRSDSIGNDRSDGTDDVGIQP   240
MicG_NODE_26_1_peptidase_partial  .......................Y......--............................   238
Op_KAH9410533.1_peptidase         .......................C......NG............................   240
```

```
Consensus                      DKQYMHDADIGTDIPHCSRVYCTHFEPSDARRMFPCFDQPDLKATFSISVDVPKTLTVLS   300
MicG_NODE_26_1_peptidase_partial ............................................................   298
Op_KAH9410533.1_peptidase      ............................................................   300

Consensus                      NTDTVPEWREDYGNRKIEYFERTCRMATYVVAIVAGSLDYVQGVSSSGIRVRVYGIAGVS   360
MicG_NODE_26_1_peptidase_partial ............................................................   358
Op_KAH9410533.1_peptidase      ............................................................   360

Consensus                      DKRSRMYALGVALRSVEYFGEYFGMRYVSCGSSAKIDMVGVPEFGAGAMENWGLVVYRTE   420
MicG_NODE_26_1_peptidase_partial ............................................................   418
Op_KAH9410533.1_peptidase      ............................................................   420

Consensus                      NLEVMAVPASAIYNGGENDREVVDEDKSMNAVDGGEIMNHNDVKNSIRDGHDKNISSENG   480
MicG_NODE_26_1_peptidase_partial ............................................................   478
Op_KAH9410533.1_peptidase      ............................................................   480

Consensus                      NGSRDSNGVFYGSSEADEDNMQQIRETVSHEIAHMWFGNLVTLRWWDELWVNEGFATWAS   540
MicG_NODE_26_1_peptidase_partial ............................................................   538
Op_KAH9410533.1_peptidase      ............................................................   540

Consensus                      LKYLQEAWGADEELRFVKDRLASSMRDEGDGRAVRDRVVSGASAAGRFDSVRYGKSASVI   600
MicG_NODE_26_1_peptidase_partial ............................................................   598
Op_KAH9410533.1_peptidase      ............................................................   600

Consensus                      RMVEQYVGPDVMREGIREYLERHKYGNVSGSDVWKAVXECWRRMKGEDSSADINDSGTSS   660
MicG_NODE_26_1_peptidase_partial .....................................G......................   658
Op_KAH9410533.1_peptidase      .....................................S......................   660

Consensus                      XSGAEISQMVREWMEQEGYPVVHVHEEDGEIVLAQSRYCSARMGVGASRVHGDTNSNSNS   720
MicG_NODE_26_1_peptidase_partial R...........................................................   718
Op_KAH9410533.1_peptidase      G...........................................................   720

Consensus                      NEEETVNEVGNSRNDNGMNGNGMDRNGDDAGCDAAKHLWRIPVSIMWDNGSMHTVLLSTP   780
MicG_NODE_26_1_peptidase_partial ............................................................   778
Op_KAH9410533.1_peptidase      ............................................................   780

Consensus                      EMRVARRTQRYKLNAGYAGYFRVHYGLNGGVLHALMVAGDSEVDQANVIEDVFELAYAGY   840
MicG_NODE_26_1_peptidase_partial ............................................................   838
Op_KAH9410533.1_peptidase      ............................................................   840

Consensus                      MSVGAVMREVLSWGTAGPWRWSVQQMQCGMETPEHSSDGNDNSNNDGLSVFARHGIEMLN   900
MicG_NODE_26_1_peptidase_partial ............................................................   898
Op_KAH9410533.1_peptidase      ............................................................   900

Consensus                      GVKTMDSTDGVNDIDGASDGMPVDKTDGXTYEIAHVPDNVVKGVVGKLLVVWGELYEDVE   960
MicG_NODE_26_1_peptidase_partial ............................R...............................   958
Op_KAH9410533.1_peptidase      ............................K...............................   960

Consensus                      VRQVLERAIWGVIGRIVKDAEICVDAVESGNETNGSSRKSDWNTKSRKGRVNGHDGDCKN  1020
MicG_NODE_26_1_peptidase_partial ............................................................  1018
Op_KAH9410533.1_peptidase      ............................................................  1020

Consensus                      RAGIEMQKYALEVGVMMGRKEAVEKARSAYERGMSRMHMPAMVGALADEXIKEMMQKYAG  1080
MicG_NODE_26_1_peptidase_partial .................................................R..........  1078
Op_KAH9410533.1_peptidase      .................................................Q..........  1080

Consensus                      AGKEWRKWVVEGMSGLMQEESVKYVLDEVCRMSRLCLYMDSKCEDSIGGCVNGLKGGKWA  1140
MicG_NODE_26_1_peptidase_partial ....--------------------------------------------------------  1138
Op_KAH9410533.1_peptidase      ............................................................  1140

Consensus                      DMNSSEMKNAEVSKEQDNVSVLSIIAGDGVINDVDATGNVMNSIVGEVLHDGGIGIADVV  1200
MicG_NODE_26_1_peptidase_partial ------------------------------------------------------------  1198
Op_KAH9410533.1_peptidase      ............................................................  1200
```

```
Consensus                          DILRGIWRGGKFRETIAEYVLRMWGAIQKAAGMNGMILRDAAEMVGGIRDVSVGVKYGDE  1260
MicG_NODE_26_1_peptidase_partial   ------------------------------------------------------------  1258
Op_KAH9410533.1_peptidase          ............................................................  1260

Consensus                          QKGKDWELAGTKVIEEIRFRERLRERRDEILSELMALFDTDGWRVLESKMHD          1312
MicG_NODE_26_1_peptidase_partial   ----------------------------------------------------          1082
Op_KAH9410533.1_peptidase          ....................................................          1312
```

**Figure S1.** Alignment of proteins with the highest number of missense mutations.

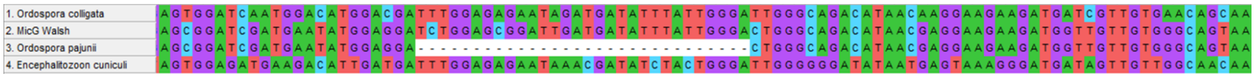

**Figure S2.** Chitin synthase gene sequence alignment of *O. colligata*, MicG, *O. pajunii* and *E. cuniculi*.

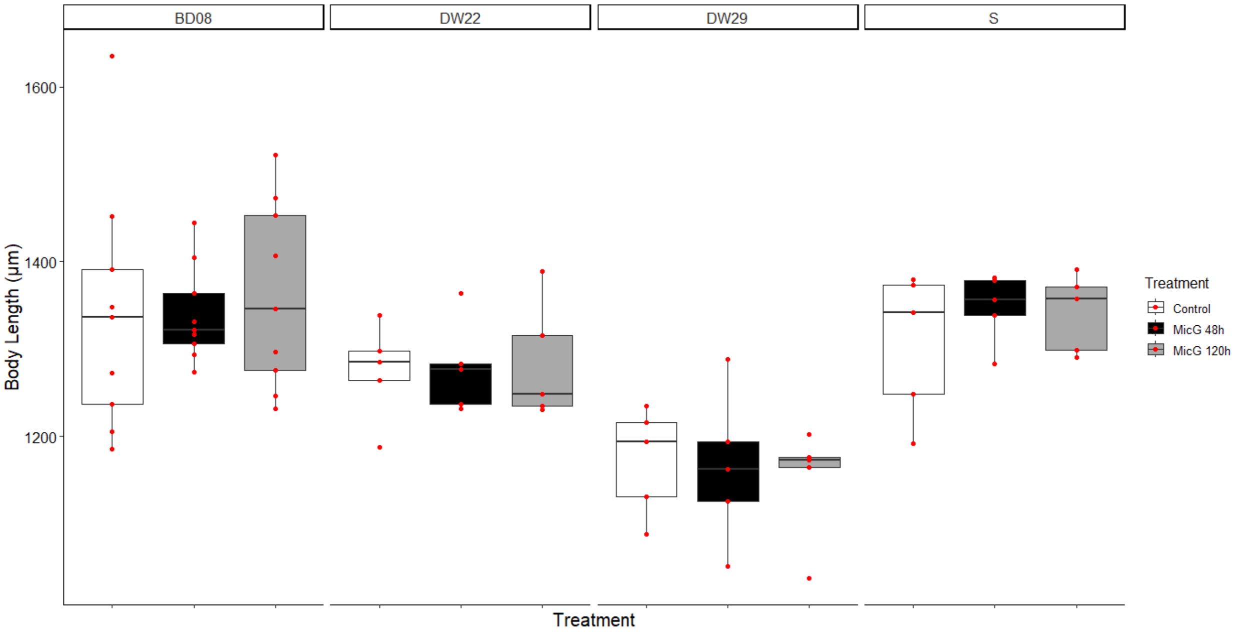

**Figure S3.** Body size of experimental *Daphnia* did not vary between MicG treatment groups.

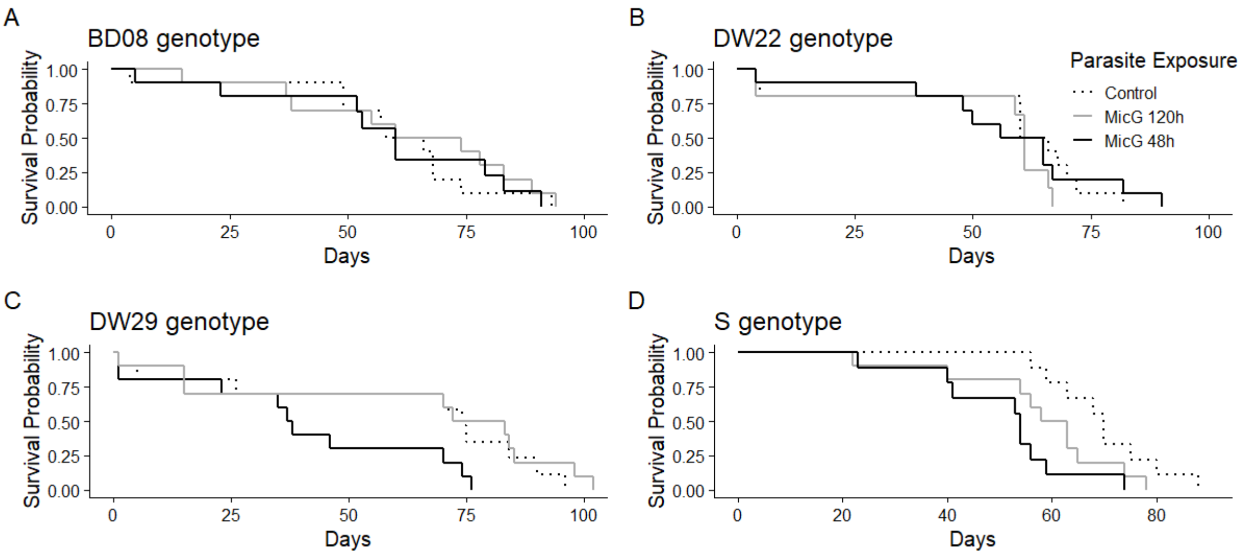

**Figure S4**. The S clone's mortality was significantly higher in MicG 48h treatment in comparison to controls (hazard ratio=4.577, 95% CI=1.616-12.964, p=0.0042). Mortality of other clones did not differ significantly between the MicG treatments and the control (Table S4).

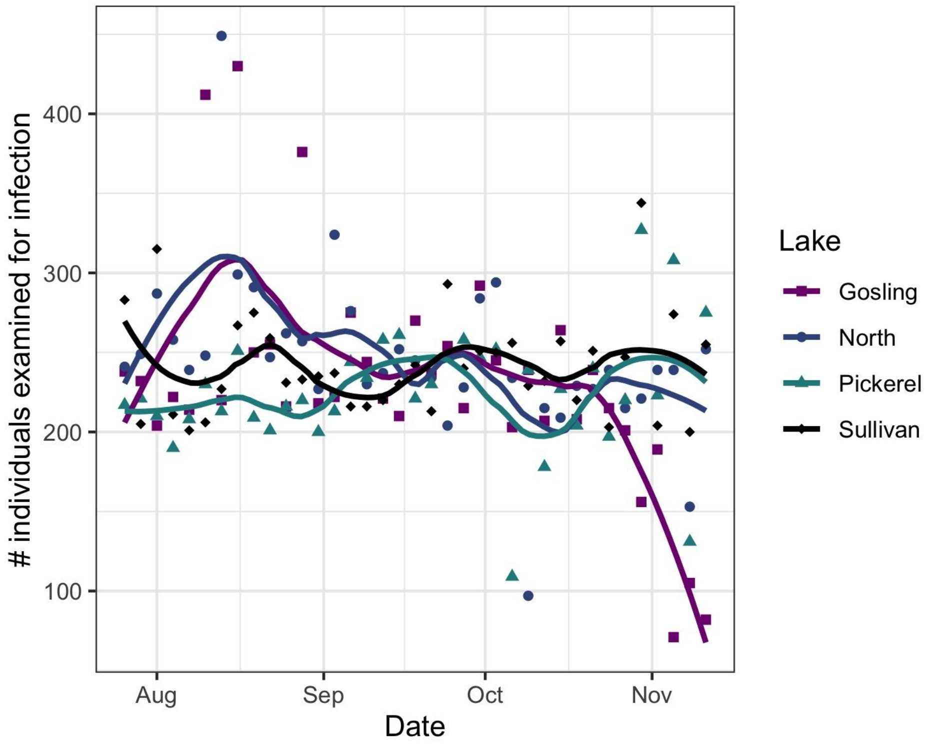

**Figure S5.** Number of *D. dentifera* individuals that were examined for infection on each date in the four study lakes.

**Table S1.** Lakes sampled from 2016-2022 that provided information on host range (all 15 lakes), on the dynamics of infection (lakes in blue), and from which samples were collected for molecular analysis (lakes in red). Walsh Lake was the source of the MicG-infected individuals that were used to start the lab cultures.

| Lake Name | Site | County | Latitude & Longitude | Max depth (m) |
|---|---|---|---|---|
| Appleton | Brighton | Livingston | 42°30'37"N, 83°50'03"W | 11.5 |
| Bishop | Brighton | Livingston | 42°30'04"N, 83°50'24"W | 16.5 |
| Bruin | Pinckney | Washtenaw | 42°25'07"N, 84°02'23"W | 14.5 |
| Cedar | Waterloo | Washtenaw | 42°18'52"N, 84°04'45"W | 7 |
| Crooked P | Pinckney | Washtenaw | 42°25'11"N, 83°58'57"W | 12 |
| Crooked W | Waterloo | Washtenaw | 42°19'32"N, 84°06'43"W | 6 |
| Gosling | Pinckney | Livingston | 42°26'22"N, 84°00'12"W | 6 |
| Little Appleton | Brighton | Livingston | 42°30'24"N, 83°50'19"W | 6 |
| Mill | Waterloo | Washtenaw | 42°19'46"N, 84°05'27"W | 6 |
| North | Pinckney | Washtenaw | 42°23'35"N, 84°00'23"W | 17 |
| Pickerel | Pinckney | Washtenaw | 42°24'37"N, 83°58'58"W | 16 |
| Sullivan | Pinckney | Washtenaw | 42°23'55"N, 84°03'25"W | 6.5 |
| Walsh | Waterloo | Washtenaw | 42°20'15"N, 84°04'47"W | 6 |
| Whitmore | Brighton | Livingston & Washtenaw | 42°25'42"N, 83°45'08"W | 19.5 |
| Woodland | Brighton | Livingston | 42°33'12"N, 83°46'29"W | 10.5 |

**Table S2.** List of genomic data used for the phylogenomic analysis

| Genus | Species | Strain | Source | Accession |
|---|---|---|---|---|
| Ordospora | pajunii | MicG | JGI | |
| Ordospora | pajunii | FI-F-10 | NCBI | GCA_021821965.1 |
| Ordospora | colligata | OC4 | NCBI | GCA_000803265.1 |
| Ordospora | colligata | GB-EP-1 | NCBI | GCA_004324935.1 |
| Ordospora | colligata | FI-SK-17-1 | NCBI | GCA_004325055.1 |
| Ordospora | colligata | NO-V-7 | NCBI | GCA_004324945.1 |
| Encephalitozoon | cuniculi | GB-M1 | NCBI | GCA_000091225.2 |
| Encephalitozoon | intestinalis | ATCC50506 | NCBI | GCA_000146465.1 |
| Encephalitozoon | romaleae | SJ-2008 | NCBI | GCA_000280035.2 |
| Encephalitozoon | hellem | ATCC50504 | NCBI | GCA_000277815.3 |
| Nosema | apis | BRL01 | NCBI | GCA_000447185.1 |
| Nosema | ceranae | BRL | NCBI | GCA_004919615.1 |
| Nosema | bombycis | CQ1 | NCBI | GCA_000383075.1 |
| Nosema | granulosis | Ou3-Ou53 | NCBI | GCA_015832245.1 |
| Enterospora | canceri | GB1 | NCBI | GCA_002087915.1 |
| Enterocytozoon | bieneusi | H348 | NCBI | GCA_000209485.1 |

| | | | | |
|---|---|---|---|---|
| Enterocytozoon | hepatopenaei | CIBAehp1 | NCBI | GCA_026692725.1 |
| Hepatospora | eriocheir | GB1 | NCBI | GCA_002087885.1 |
| Vittaforma | corneae | ATCC50505 | NCBI | GCA_000231115.1 |
| Anncaliia | algerae | PRA339 | NCBI | GCA_000385875.2 |
| Edhazardia | aedis | USNM41457 | NCBI | GCA_000230595.3 |
| Tubulinosema | ratisbonensis | Franzen | NCBI | GCA_004000155.1 |
| Thelohania | contejeani | T1 | NCBI | GCA_014805555.1 |
| Hamiltosporidium | magnivora | BE-OM-2 | NCBI | GCA_004325065.1 |
| Hamiltosporidium | tvaerminnensis | FI-OER-3-3 | NCBI | GCA_022605425.1 |
| Cucumispora | dikerogammari | Dv6 | NCBI | GCA_014805705.1 |
| Spraguea | lophii | Celtic_Deep | NCBI | GCA_001887945.1 |
| Trachipleistophora | hominis | | NCBI | GCA_000316135.1 |
| Vavraia | culicis | floridensis | NCBI | GCA_000192795.1 |
| Pseudoloma | neurophilia | MK1 | NCBI | GCA_001432165.1 |
| Nematocida | displodere | JUm2807 | NCBI | GCA_001642395.1 |
| Nematocida | parisii | ERTm1 | NCBI | GCA_000250985.1 |
| Nematocida | ausubeli | ERTm6 | NCBI | GCA_000738915.1 |
| Nematocida | ironsii | ERTm5 | NCBI | GCA_001642415.1 |

| | | | | |
|---|---|---|---|---|
| Dictyocoela | roeselum | Ou19 | NCBI | GCA_016255985.1 |
| Dictyocoela | muelleri | Ou54 | NCBI | GCA_016256075.1 |
| Antonospora | locustae | CLX | NCBI | GCA_007674295.1 |
| Amphiamblys | sp. | WSBS2006 | NCBI | GCA_001875675.1 |
| Metchnikovella | incurvata | LNA5 | NCBI | GCA_003600395.1 |

**Table S3.** Proportion of clone S and *D. longispina* clones displaying visible infection. 10 individuals were exposed for each of the clones.

| Genotype | Proportion Infected |
|---|---|
| S | 1 |
| OLCH 17 | 0 |
| STEC 8 | 0 |
| STEC 49 | 0.8 |
| WALD 12 | 0 |
| WALD 37 | 0.9 |

**Table S4.** Statistical analysis of survival in all experimental clones, including full model (with three treatments: control, MicG 48h, and MicG 120h) and pairwise comparisons of MicG 48h and MicG 120h to the control treatment.

| Genotypes | Full Model | | | Control:MicG 48h | | | Control:MicG 120h | | |
|---|---|---|---|---|---|---|---|---|---|
| | ChiSq | DF | P | HR | 95%CI | P | HR | 95%CI | P |
| S | 8.415 | 2 | 0.015 | 4.577 | 1.616-12.964 | 0.042 | 2.267 | 0.8553-6.009 | 0.01 |
| BD08 | 0.84 | 2 | 0.657 | 1.039 | 0.425-2.539 | 0.933 | 0.709 | 0.285-1.769 | 0.461 |
| DW22 | 1.385 | 2 | 0.5 | 0.916 | 0.37-2.269 | 0.85 | 1.66 | 0.616-4.467 | 0.316 |
| DW29 | 9.58 | 2 | 0.008 | 2.717 | 0.955-7.731 | 0.061 | 0.428 | 0.143-1.28 | 0.13 |
